# Dendritic inhibition terminates plateau potentials in CA1 pyramidal neurons

**DOI:** 10.1101/2025.06.05.657434

**Authors:** Lee O. Vaasjo, Shawn Kotermanski, Tiya Patel, Hengyue J. Shi, Robert Machold, Simon Chamberland

## Abstract

In CA1 pyramidal neurons (CA1-PYRs), plateau potentials control synaptic plasticity and the emergence of place cell identity. Here, we show that dendritic inhibition terminates plateaus in an all-or-none manner. Plateaus were initially resistant to inhibition but became increasingly susceptible to termination as they progressed. Between two distinct subtypes of dendrite-targeting interneurons, OLM^Ndnf^ generated slower postsynaptic currents that terminated plateaus more effectively than OLM^α2^. Voltage-gated Ca^2+^ channels (VGCCs) were necessary for plateaus, which were prolonged by blocking small-conductance Ca^2+^-activated K^+^ channels (SK). A single-compartment model with these two conductances recapitulated core experimental findings and provided a mechanistic explanation for terminations. Plateaus arose from VGCCs maintained in the active state by sustained Ca^2+^ influx, a positive feedback loop that was quasi-balanced by I_SK_. Inhibition terminated plateaus by driving the membrane potential below a dynamic threshold to deactivate VGCCs and end the positive feedback loop. Lastly, two-photon Ca^2+^ imaging showed that plateaus evoke large dendritic Ca^2+^ transients that were graded by terminations. Overall, our results demonstrate how the feedback inhibitory circuit interacts with intrinsic cellular mechanisms to regulate plateau potentials and shape dendritic Ca^2+^ signals in CA1-PYRs.

## Introduction

Neurons actively integrate information in their dendrites where multiple forms of dendritic spikes amplify synaptic potentials. Plateau potentials are a form of dendritic spikes characterized by prolonged membrane depolarization. In hippocampal CA1 pyramidal cells (CA1-PYRs), plateaus evoked in single dendritic branches can last for hundreds of milliseconds and contribute to multiple forms of synaptic plasticity (1–4). In behavioral timescale plasticity (BTSP), a form of one-shot learning, a single plateau suffices to strengthen excitatory synaptic inputs to enable place field formation in CA1-PYRs (5). Therefore, plateaus are a biophysical signal controlling synaptic plasticity and memory.

Inhibition and membrane hyperpolarization interfere with multiple forms of dendritic spikes including plateaus (6–16). Plateaus present a depolarized state with elevated driving force for GABA_A_R-mediated Cl^-^ currents. In CA1-PYRs, plateaus are generated and maintained by multiple conductances, likely much smaller than those supporting action potentials, potentially rendering plateaus vulnerable to synaptic activity (1, 3, 4, 17, 18). How inhibition controls ongoing plateaus in CA1-PYRs remains poorly understood.

A heterogeneous population of GABAergic interneurons (INs) controls activity across specific spatial domains of CA1-PYRs (13, 15, 19–21). Among these, somatostatin-expressing interneurons (*Sst*-INs) provide key feedback inhibition to CA1-PYRs dendrites (3, 4, 22–25), shaping local synaptic input integration (26–29), burst firing (30), and hippocampal-dependent memory (31–34). Single-cell transcriptomics and intersectional genetic tools have recently confirmed extensive heterogeneity within *Sst*-INs and enabled access to distinct subtypes (35, 36). We recently identified that neurons co-expressing *Ndnf* and *Nkx2-1* adopt an OLM identity (OLM^Ndnf^) and preferentially innervate CA1-PYRs, in contrast to *Chrna2*-expressing OLMs (OLM^α2^) (28, 35). How subtypes of OLMs control CA1-PYRs dendritic activity remain unknown.

Here, we discovered that dendritic inhibition terminates plateaus in an all-or-none manner in CA1-PYRs. Plateaus were initially resistant to inhibitory termination but became increasingly susceptible as they progressed over time. Optogenetic stimulation of OLM^Ndnf^ terminated plateaus more effectively than OLM^α2^, an effect attributed to the slower synaptic currents mediated by OLM^Ndnf^. Experiments and biophysical modelling revealed the mechanisms governing plateau termination, highlighting that synaptic inhibition must overcome rebalancing of intrinsic conductances to drive V_M_ below a dynamic threshold. Finally, two-photon Ca^2+^ imaging revealed that plateau termination effectively grades dendritic Ca^2+^ elevations.

## Results

### Dendritic inhibition terminates plateaus in CA1-PYRs

Plateau potentials are prolonged depolarizations in CA1-PYRs that control synaptic plasticity and memory formation (1, 2, 5, 37). *Sst*-INs primarily synapse onto CA1-PYRs dendrites where they modulate local activity (21, 24, 26, 35, 38). How *Sst*-INs control ongoing plateaus in CA1-PYRs has yet to be elucidated.

We performed whole-cell recordings from deep CA1-PYRs with Cs^+^-based intracellular solution in acute hippocampal slices from *Sst-Cre;;Ai32* mice which express channelrhodopsin-2(H134R) in *Sst*-INs (Fig.1 A) (39). Current injection (50 ms, 50 – 200 pA, 118.2 ± 4.6 pA, n = 51) generated a single or few action potentials (APs) followed by plateaus of long duration (140.2 ± 10.2 ms, n = 51, Fig. 1B). Optogenetic activation of *Sst*-INs with a brief pulse of blue light (2 ms, 450 nm) terminated plateaus in an all-or-none manner (Fig. 1B). Light power was adjusted at the threshold value to observe terminations (0.7 ± 0.2 mW, 44.7 ± 6.3%, n = 11) and doubling the light power significantly increased the likelihood of plateau termination (1.4 ± 0.4 mW, 92.7 ± 3.4%, n = 11, p < 0.001, Student’s paired t-test; Fig. 1C – D). In cases where optogenetic stimulation failed to terminate plateaus, an inhibitory postsynaptic potential (IPSP) was visible and the V_M_ returned to the depolarized phase where the plateau pursued its normal course as if the IPSP-induced disturbance had no lasting impact (plateau duration without IPSP = 162.4 ± 20.1 ms vs. with IPSP = 180.9 ± 15.9 ms, n = 14, p = 0.15, Wilcoxon signed-rank test, Fig. 1E; break point without IPSP = −9.9 ± 1 mV vs. with IPSP = −9.3 ± 1 mV, n = 14, p = 0.33, Student’s paired t-test, Fig. 1G). Inspection of individual traces revealed that terminations happened around a similar point in V_M_ suggesting that a putative threshold had to be reached (Fig. 1C, E). These results show that *Sst*-INs terminate plateaus in an all-or-none manner in CA1-PYRs.

**Figure 1.**
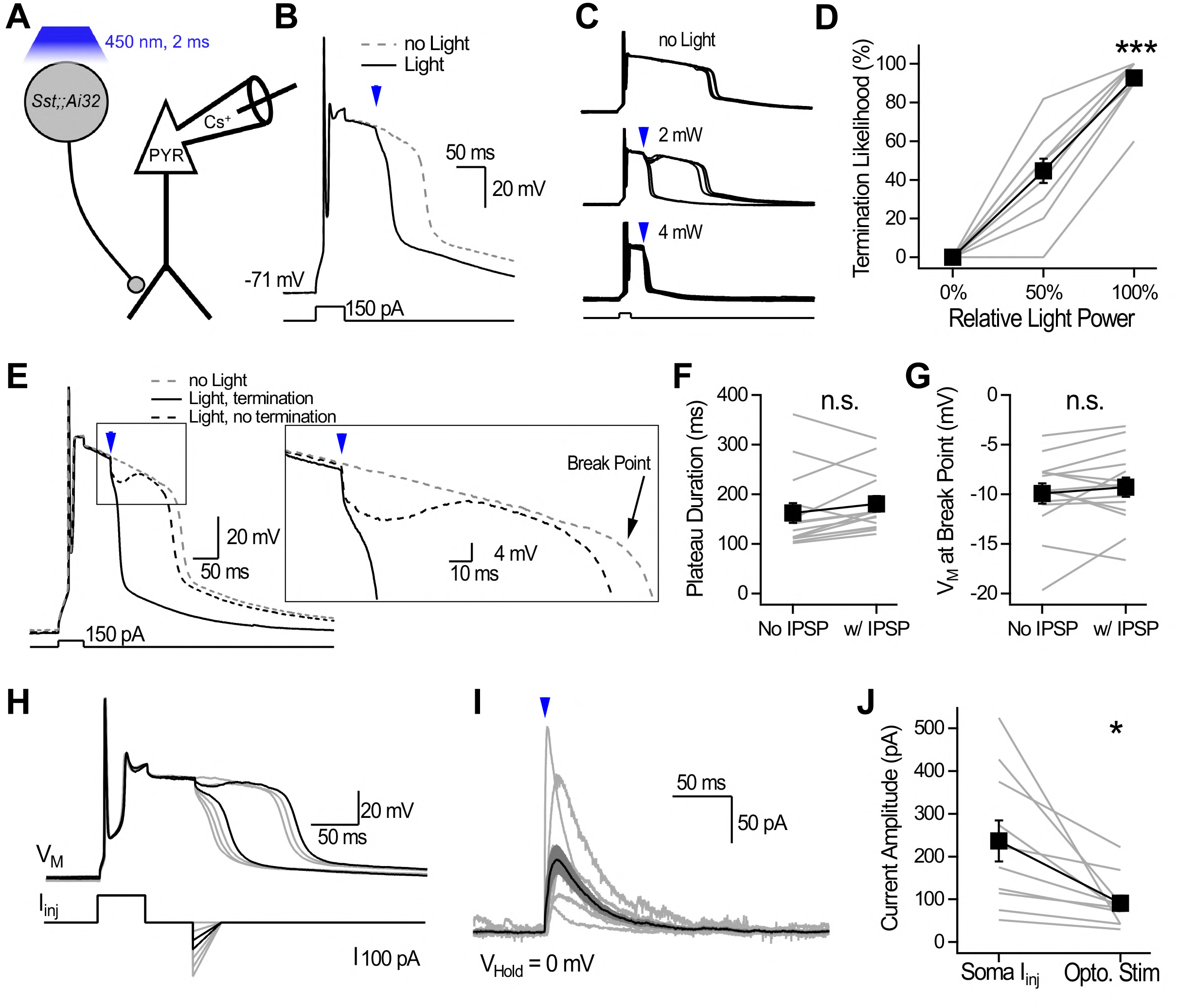
Optogenetic activation of *Sst*-INs terminates plateaus in an all-or-none manner in CA1-PYRs. **A**, Scheme showing the recording configuration. **B**, Current-clamp recording from a CA1-PYR. Brief current injections evoked long-lasting plateaus that were terminated by optogenetic stimulation of *Sst*-INs. **C**, Current-clamp recordings from a CA1-PYR showing plateaus in absence of light (top), and with application of blue light (middle, 2 mW; bottom, 4 mW). 5 consecutive trials are shown for each condition. **D**, Termination likelihood as a function of light power. **E**, Traces showing plateaus in absence of light and with sub- and suprathreshold light. An IPSP is evident for subthreshold optogenetic stimulation. **F**, Plateau duration in absence of light (No IPSP) and with a subthreshold IPSP (w/ IPSP). **G**, Membrane potential at natural break point for plateaus in absence of light and with a subthreshold IPSP. **H**, Current injection mimicking IPSPs terminate plateaus. Example shows 7 consecutive traces for which the mock IPSP amplitude was gradually increased. Black traces show the just sub- and suprathreshold conditions. **I**, Voltage-clamp recording (0 mV) showing IPSCs evoked by optogenetic stimulation. Power was adjusted at the just-threshold value to observe both terminations and failures. Gray traces show individual neuron average, and the black trace shows the population average ± SEM (n = 10 neurons). **J**, Current amplitude required for just threshold terminations when delivered at the soma compared to the IPSC amplitude recorded at the soma for just threshold terminations evoked optogenetically. For D, F, G and J: gray lines show individual neurons, and the black trace shows average ± SEM. n.s. = non-significant; * p < 0.05; ** p < 0.01; and *** p < 0.001

Optogenetic stimulation caused IPSPs that invariably preceded plateau termination (Fig. 1B, C, E). We tested whether membrane hyperpolarization suffices to terminate plateaus by injecting IPSP-mimicking current waveforms via the recording electrode. Mock IPSPs were approximated by an instantaneous hyperpolarizing current of increasing amplitude that decayed over 30 ms (Fig. 1H). Mock IPSPs terminated plateaus in an all-or-none manner as a function of current amplitude (n = 14/14 neurons tested, Fig. 1H – I). The minimal somatic current injection required to terminate plateaus was higher than the optogenetically-evoked IPSCs (V_Hold_ = 0 mV) that were associated with just-threshold plateau terminations (soma injection: 236.6 ± 48.2 pA; optogenetic stimulation: 90.6 ± 18 pA, n = 10, p < 0.05, Student’s paired t-test, Fig. 1J), indicating that plateaus are more easily terminated by *Sst*-INs than somatic current injection. Overall, these results show that while inhibition provides the necessary membrane hyperpolarization, plateau termination is operated by a cell-autonomous mechanism.

### Plateaus become progressively more susceptible to termination

The duration of plateaus recorded in CA1-PYRs is variable (40, 41). Excitatory synapses formed by CA1-PYRs on *Sst*-INs demonstrate modest quantal size but show considerable short-term facilitation, suggesting that *Sst*-INs could be optimally recruited during CA1-PYRs repetitive firing (22, 23, 35, 42, 43). Consequently, dendritic inhibition mediated by *Sst*-INs could occur at multiple time points during plateaus.

Optogenetic stimulation of *Sst*-INs was delivered 50 ms following plateau induction and the light power was adjusted near the termination threshold in each cell (light power: 3.9 ± 1.3 mW; termination likelihood: 62.7 ± 7.7%; n = 17, Fig. 2A – B). Identical optogenetic stimulation delivered at 10, 20, 50 and 100 ms during the plateau revealed a significant effect of stimulation timing on termination likelihood (Friedman test: χ^2^ = 42.8, p < 10^-8^), with earlier illumination being significantly less effective at terminating plateaus (10 ms: 20 ± 6.7%, p < 0.01; 20 ms: 33.3 ± 8.8%, p < 0.01; both compared to 50 ms: 62.7 ± 7.7%, n = 17, post-hoc Wilcoxon signed-rank tests with Holm-Bonferroni correction; Fig. 2B). These results show that plateaus are initially resistant but become increasingly susceptible to termination.

**Figure 2.**
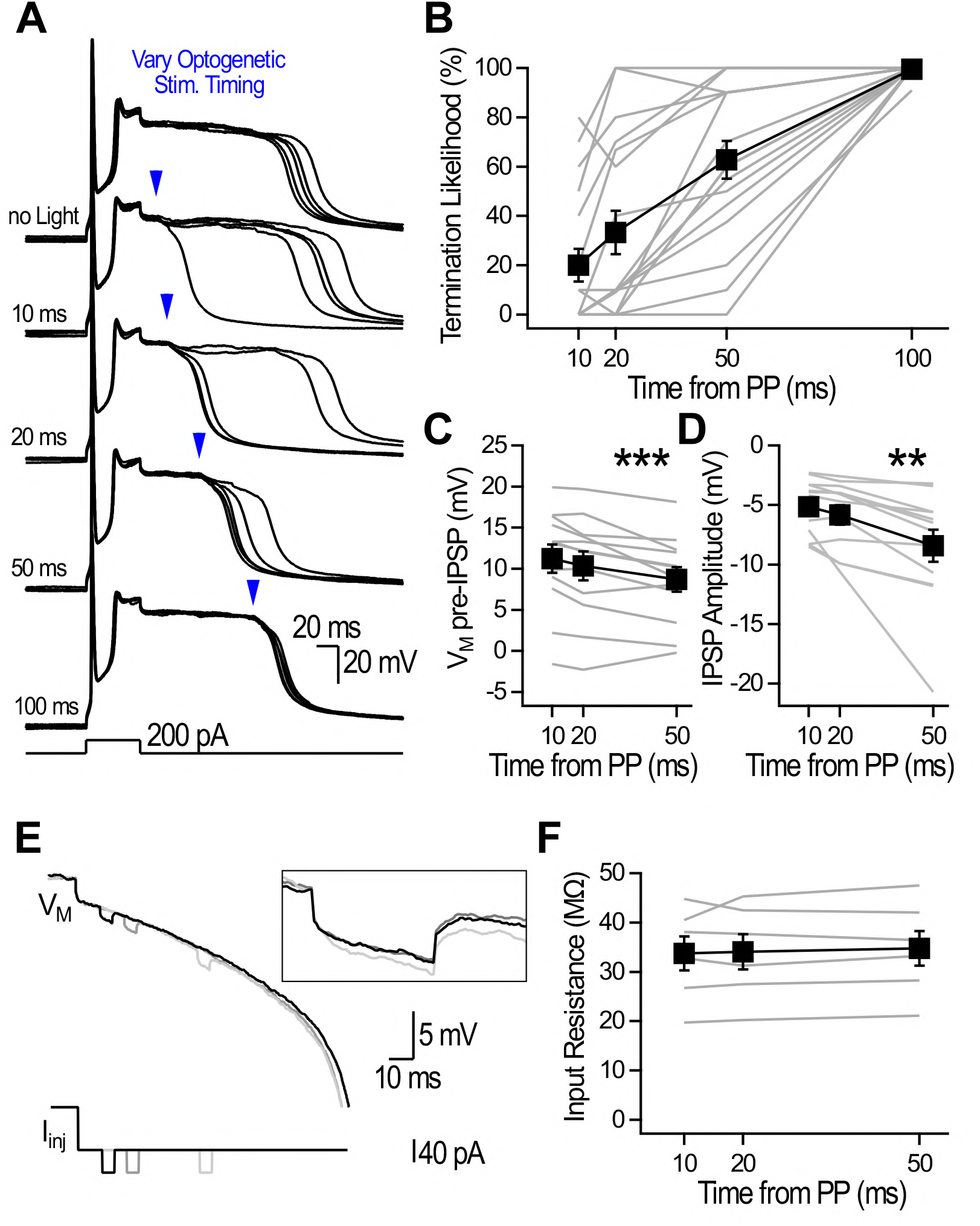
Plateaus are initially resistant but become gradually more sensitive to inhibition. **A**, Plateaus without light application (top) and with optogenetic stimulation delivered at 4 timepoints during plateaus. Examples shown are 5 consecutive traces for each condition. Light power was adjusted at 50 ms to obtain both success and failures and was kept constant for all measurements within a neuron. **B**, Termination likelihood as a function of optogenetic timing. **C**, Membrane potential just before the IPSP onset for 10, 20 and 50 ms timepoints. **D**, IPSP amplitude for subthreshold IPSPs evoked at 10, 20, and 50 ms timepoints. **E**, Brief current injection via the recording pipette at 10, 20 and 50 ms following plateau induction. There was no difference in the input resistance evolution whether the injected current was depolarizing or hyperpolarizing, and results were pooled. **F**, The measured input resistance did not change as a function of current injection timing. ** p < 0.01; and *** p < 0.001

We next investigated why early resistance to inhibition gradually dissipates. We analyzed trials in which plateaus were not terminated and measured: 1) the V_M_ right before optogenetic stimulation, and 2) the light-evoked IPSP amplitude as a function of time from plateau onset. A significant hyperpolarizing shift in V_M_ during plateaus (n = 12, Fig. 2C) was paralleled by gradually larger IPSP amplitudes (n = 12, Fig. 2D). The observed increase in IPSP amplitude is inconsistent with the decrease in Cl^-^ driving force caused by the gradual hyperpolarizing shift in V_M_, prompting us to examine the evolution of membrane resistance. Brief current steps delivered via the recording pipette at corresponding timepoints revealed that the input resistance remained constant during plateaus (repeated-measures ANOVA: F(2,10) = 0.017, p = 0.98; Fig. 2E). Overall, these results indicate that feedback inhibition governs plateau duration, with progressive V_M_ hyperpolarization and increasing IPSP amplitude both enhancing susceptibility to termination over time.

### OLM^Ndnf^ preferentially terminate plateaus compared to OLM^α2^

*Sst*-INs are genetically, anatomically, and functionally diverse (19, 35, 44, 45). A proportion of *Sst*-INs located in stratum oriens project their axon to stratum lacunosum-moleculare (OLMs) (24, 46–49). We recently identified that the *Ndnf;;Nkx2-1* genetic intersection provides access to OLM^Ndnf^, a subpopulation which preferentially innervate CA1-PYRs, unlike previously described OLM^α2^ (28, 35).

To investigate how OLM subtypes control plateaus, we generated *Ndnf-FlpO;;Nkx2-1-Cre;;Ai80* and *Chrna2-Cre;;Sst-FlpO;;Ai80* mice. Optogenetic stimulation was kept constant across all experiments (10 mW, 2 ms) and delivered 10, 20, 50 and 100 ms after plateau initiation (Fig. 3A – C). As expected, termination likelihood for both OLM^Ndnf^ and OLM^α2^ increased as a function of time during plateaus (Fig. 3C). Interestingly, termination likelihood was higher for OLM^Ndnf^ than OLM^α2^, an effect that was significant at the 10, 20 and 50 ms timepoints (for the 50 ms timepoint: OLM^Ndnf^: 84.4 ± 9.6%, n = 11; OLM^α2^: 42.6 ± 9.4%; n = 16, p < 0.01, Mann Whitney U test, Fig. 3C). To investigate why OLM^Ndnf^ were more likely to terminate plateaus, we recorded inhibitory postsynaptic currents (IPSCs) in CA1-PYRs voltage-clamped at 0 mV (Fig. 3D – E). Optogenetic stimulation of OLM^Ndnf^ and OLM^α2^ evoked IPSCs of similar amplitude and rise time (amplitude: OLM^Ndnf^: 105.9 ± 10.7 pA, n = 23; OLM^α2^: 103.4 ± 13.3 pA, n = 22, p = 0.6, Mann Whitney U test, Fig. 3F; rise time: OLM^Ndnf^: 7.6 ± 0.5 ms, n = 23; OLM^α2^: 6.4 ± 0.3 ms, n = 22, p = 0.1, Mann Whitney U test, Fig 3G). However, the decay kinetics of IPSCs evoked by OLM^Ndnf^ photostimulation were significantly slower than those evoked by identical photostimulation of OLM^α2^ (decay τ: OLM^Ndnf^: 46.5 ± 2.4 ms, n = 23; OLM^α2^: 35.4 ± 1.5 ms, n = 22, p < 0.001, unpaired Student’s t-test, Fig. 3H). Overall, these results show that OLM^Ndnf^ and OLM^α2^ can terminate plateaus, but OLM^Ndnf^ do so more effectively.

**Figure 3.**
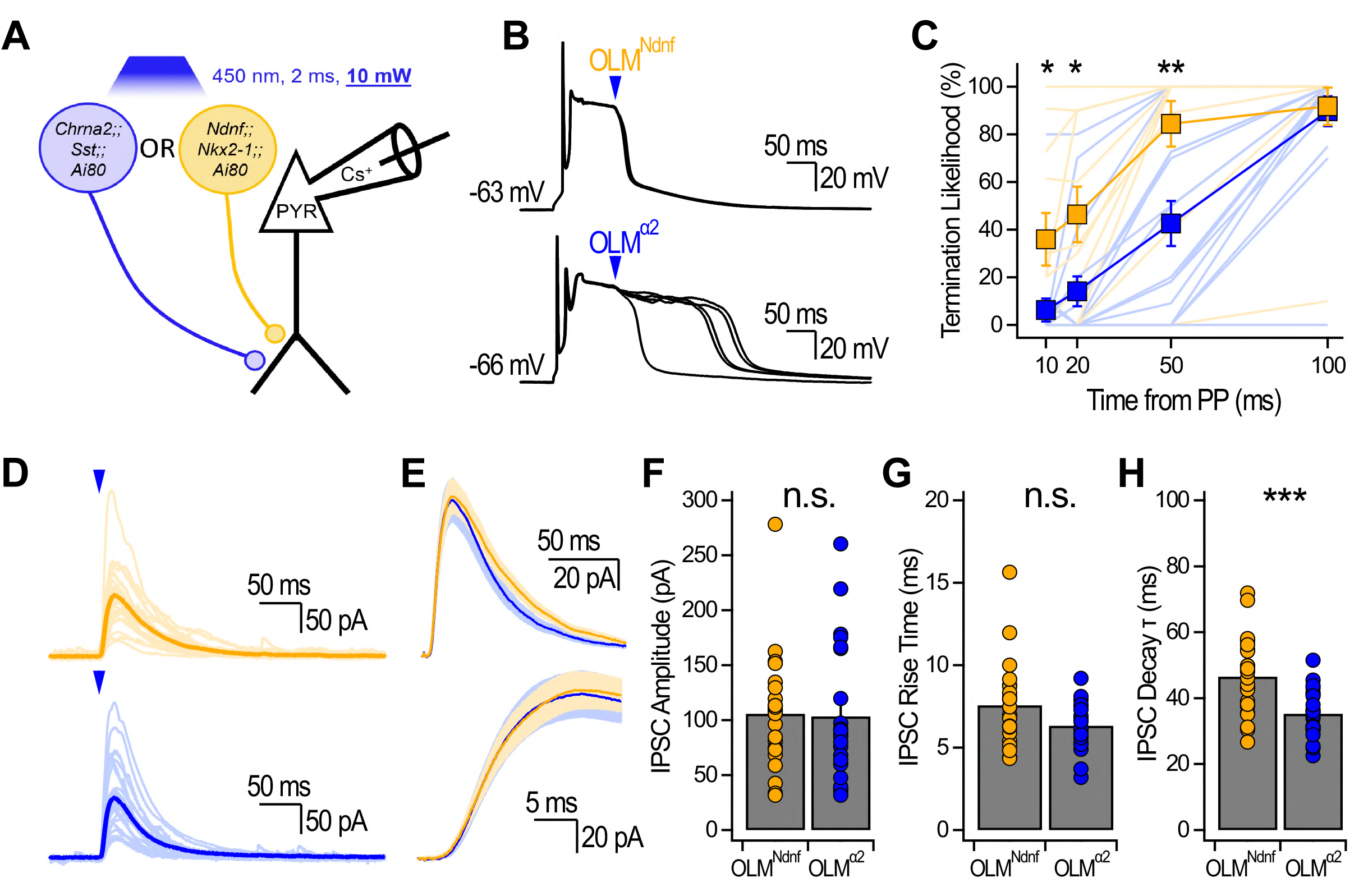
OLM^Ndnf^ terminate plateaus more effectively than OLM^α2^. **A**, Scheme showing the recording configuration. The light power was kept constant at 10 mW across all CA1-PYRs recorded in both transgenic mouse models. **B**, Example traces showing the impact of optogenetic stimulation of OLM^Ndnf^ and OLM^α2^. 5 consecutive traces are shown. **C**, Termination likelihood as a function of optogenetic stimulation timing. For the 10 ms timepoint: OLM^Ndnf^: 35.9 ± 11.1%, n = 11; OLM^α2^: 6.3 ± 4.8%; n = 16, p < 0.05, Mann Whitney U test. For the 20 ms timepoint: OLM^Ndnf^: 46.4 ± 11.7%, n = 11; OLM^α2^: 14.2 ± 6.3%; n = 16, p < 0.05, Mann Whitney U test. **D**, IPSCs recorded in voltage-clamp (0 mV) evoked by constant optogenetic stimulation. Light color traces show the average for individual neurons. Population averages are shown in darker traces with shaded area showing ± SEM. **E**, Population averages at two timescales to show differences in decay and in rise time. Shaded areas show ± SEM. **F**, Optogenetically-evoked IPSCs amplitude recorded for OLM^Ndnf^ and OLM^α2^. **G**, IPSC rise time (20 – 80% of maximum amplitude) for both OLM subtypes. **H**, IPSC decay τ for both OLM subtypes. n.s. = non-significant; * p < 0.05; ** p < 0.01 and *** p < 0.001.

Functional differences prompted us to more closely evaluate the identity of OLM^Ndnf^ and OLM^α2^ which show transcriptomic proximity and adopt a common anatomical phenotype (35, 36). To test how this translates *in situ* with the transgenic mouse models, we generated triple transgenic *Ndnf-FlpO;;Chrna2-Cre;;Ai224* mice to drive the nuclear expression of EGFP and dTomato as a function of Cre- and FlpO-expression, respectively (50) (Fig. 4A – B). Analysis of confocal images revealed that EGFP and dTomato expression was distributed as expected from previous data in *Chrna2*-Cre mice (28) and *Ndnf* transcripts in the hippocampus (51) (Fig. 4A – B). We found that only 8.3% of GFP-expressing cells in stratum oriens co-expressed dTomato (Fig. 4C). We chose this quantification approach because *Ndnf* is expressed in multiple interneuron subtypes outside of the *Nkx2-1* intersection that targets OLM^Ndnf^, while *Chrna2* on its own targets OLM^α2^ (28, 35). Minimal colocalization between the two fluorescent proteins suggested that OLM^Ndnf^ and OLM^α2^ are nearly non-overlapping subpopulations. We next evaluated potential differences in axonal projections that could help account for the distinct IPSCs evoked by OLM^Ndnf^ and OLM^α2^ with biocytin fills of genetically targeted neurons followed by Neurolucida reconstructions (Figs. 4D, S1 and S2). Axonal distribution analysis through the CA1 layers revealed that OLM^α2^ axons penetrated deeper in LM than those of OLM^Ndnf^ (Fig. 4E). Overall, OLM^Ndnf^ and OLM^α2^ are distinct subpopulations of OLMs that terminate plateaus, but OLM^Ndnf^ do so more effectively. OLM^Ndnf^ mediate slower IPSCs, a finding that is unlikely to be explained by differences in electrotonic attenuation along CA1-PYR dendrites.

**Figure 4.**
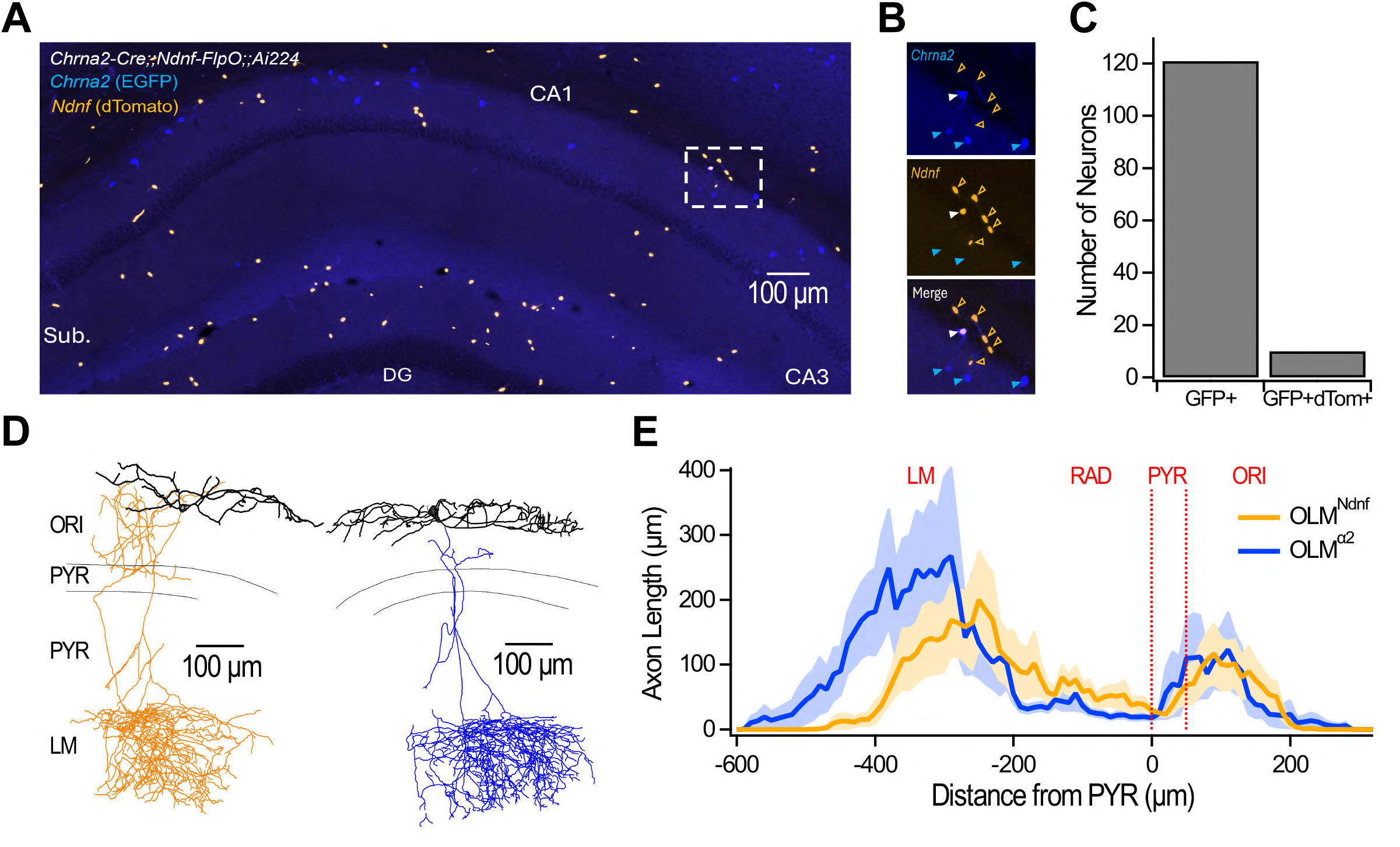
OLM^Ndnf^ and OLM^α2^ minimally overlap and possess distinct axonal arborizations. **A**, Confocal image obtained from a hippocampal section (50 µm) of a triple transgenic *Chrna2-Cre;;Ndnf-FlpO;;Ai224* mouse. **B**, Insets show examples of overlapping and non-overlapping neurons. Note that while dTomato is restricted to the soma, GFP expression is also seen in the cytosol as reported before for this reporter line (50). **C**, Quantification of overlapping and non-overlapping neurons sampled across multiple images. Because *Ndnf*-expressing neurons in stratum oriens and in the alveus could be from a different subtype (such as neurogliaform), we chose to report the number of GFP cells that co-express dTomato. **D**, Neurolucida reconstructions of an OLM^Ndnf^ and an OLM^α2^ recorded in *Ndnf-FlpO;;Nkx2-1-Cre;;Ai65* and *Chrna2-Cre;;Ai9* transgenic animals, respectively. **E**, Axonal length as a function of the distance from the pyramidal cell layer. Dark lines show the average across all cells, and shaded areas show SEM. All neurons reconstructed and reported here are shown in Figs. S1 and S2.

### Plateaus require voltage-gated Ca^2+^ channels but not voltage-gated Na^+^ channels

We sought to dissect the biophysical mechanisms underlying plateau termination through experiments and modeling. The constant input resistance observed during plateaus argues that plateaus are unlikely to originate from a single conductance that gradually decays over time but instead reflects a near-equilibrium between multiple conductances (Fig. 2E). Dendritic plateaus are abolished by application of voltage-gated Ca^2+^ channels blockers and prolonged by SK channel blockade (3, 4, 17). In addition, TTX-sensitive Na^+^ channels have been shown to contribute to multiple forms of regenerative activity in CA1-PYR dendrites (1, 4, 52, 53). We next pharmacologically dissected the roles of these currents in plateaus and their termination.

Bath application of the Na^+^ channel blocker TTX (1 µM) blocked APs and plateau generation in all neurons tested (n = 32). Increasing the current injection amplitude (control: 110.9 ± 5.2 pA; TTX: 296.1 ± 12.5 pA, n = 32, p < 0.001, Wilcoxon signed-rank test, Fig. 5A – B) consistently rescued plateaus without restoring APs (plateau likelihood: control: 99.5 ± 0.4%; TTX: 0%; increased current injection in TTX: 98.7 ± 0.8%, n = 31; Fig. 5A – B). Plateaus evoked in TTX were significantly briefer than in control condition (control: 132.4 ± 13 ms; TTX: 81.3 ± 10.2 ms; n = 32, p < 0.001, Wilcoxon signed-rank test, Fig. 5B), indicating that while TTX-sensitive Na^+^ currents are not essential for plateau generation, they likely enhance local depolarization.

**Figure 5.**
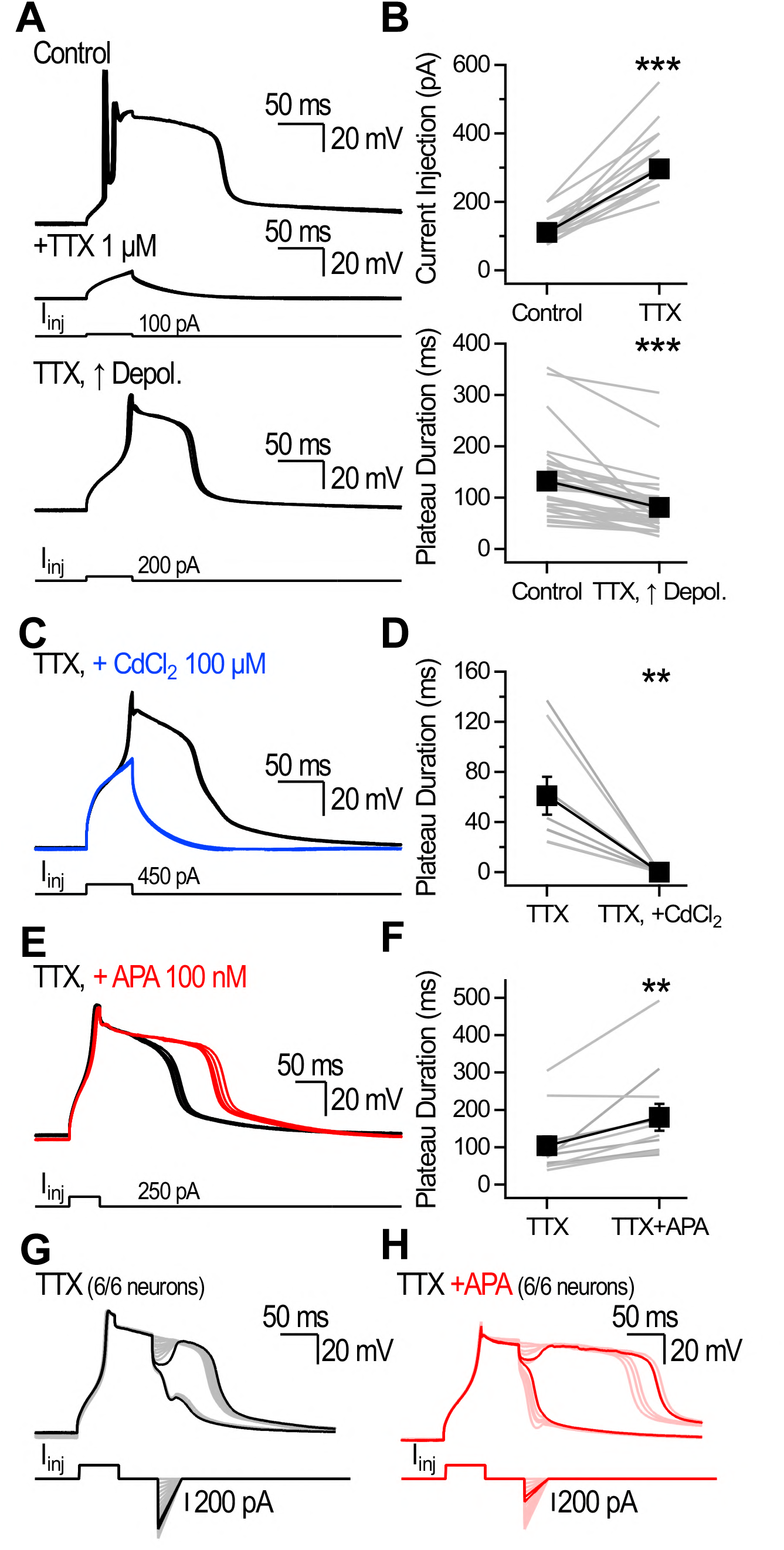
Pharmacological dissection of plateaus. **A**, Plateaus recorded in control (top) and in the presence of TTX (1 µM). Action potentials and plateaus were initially abolished (middle). Increasing the current injection amplitude recovered the plateaus but not the action potentials (bottom). Examples show 5 consecutive traces. **B**, Just-suprathreshold current injection for neurons recorded in control and following bath application of TTX (top). Plateau duration in control and following recovery by increased depolarization in TTX (bottom). **C**, Plateaus recorded in the presence of TTX (black) and following the addition of CdCl_2_ (100 µM, blue). 5 consecutive traces are shown for both. **D**, Plateau duration recorded in the presence of TTX and in the presence of TTX + CdCl_2_. **E**, Plateaus recorded in the presence of TTX (black) and following perfusion of TTX + apamin (APA, 100 nM, red). **F**, Plateau duration measured in TTX and following the application of TTX + APA. **G**, **H**, Plateaus were terminated by mock IPSPs in the presence of TTX (6/6 neurons) and in the presence of TTX + apamin (6/6 neurons). Examples show successive sweeps in which the mock IPSP amplitude was gradually increased. Dark lines show the just-subthreshold and just-suprathreshold trials for the V_M_ recordings and the injected current.

Application of Cd^2+^ (100 µM) following TTX abolished plateaus (n = 8, Fig. 5C – D). Increasing the injected current (333.3 ± 28.1 pA) by more than three-fold (1058.3 ± 96.8 pA) failed to rescue plateaus (n = 6, Fig. 5C – D), consistent with the necessity of VGCCs. Application of the SK channel blocker apamin (APA, 100 nM) after TTX significantly extended plateau duration (TTX: 104.1 ± 24.9 ms; TTX + APA: 179.7 ± 36.2 ms, n = 11, p < 0.01, Wilcoxon signed-rank test, Fig. 5E – F). Importantly, mock-IPSPs terminated plateaus in the presence of TTX (n = 6/6 cells tested, Fig. 5G) and in TTX + APA (n = 6/6 cells tested, Fig. 5H). These results show that although TTX-sensitive and apamin-sensitive currents modulate plateau dynamics, they are not required for IPSP-driven plateau termination. Overall, these results point to the central role of VGCCs in plateaus and their termination.

### Biophysical modeling provides a mechanistic explanation for plateau termination

We next aimed to understand the biophysical mechanisms controlling plateau termination. The combination of VGCCs and SK currents (I_VGCC_ and I_SK_, respectively) with opposite polarity could support a depolarized state. A single-compartment model with VGCCs and SK conductances generated plateaus that outlasted the current injection before spontaneously repolarizing to resting V_M_ (Fig. 6A and Fig. S3). The model captured key experimental findings: 1) IPSPs terminated plateaus in an all-or-none manner (Fig. 6A1 – A2 and Fig. 1); 2) plateaus were initially resistant to inhibition but became increasingly susceptible to termination over time (Fig. 6B1 – B2 and Fig. 2); 3) slow IPSPs terminated plateaus more effectively (Fig. 6C1 – C2 and Fig. 3).

**Figure 6.**
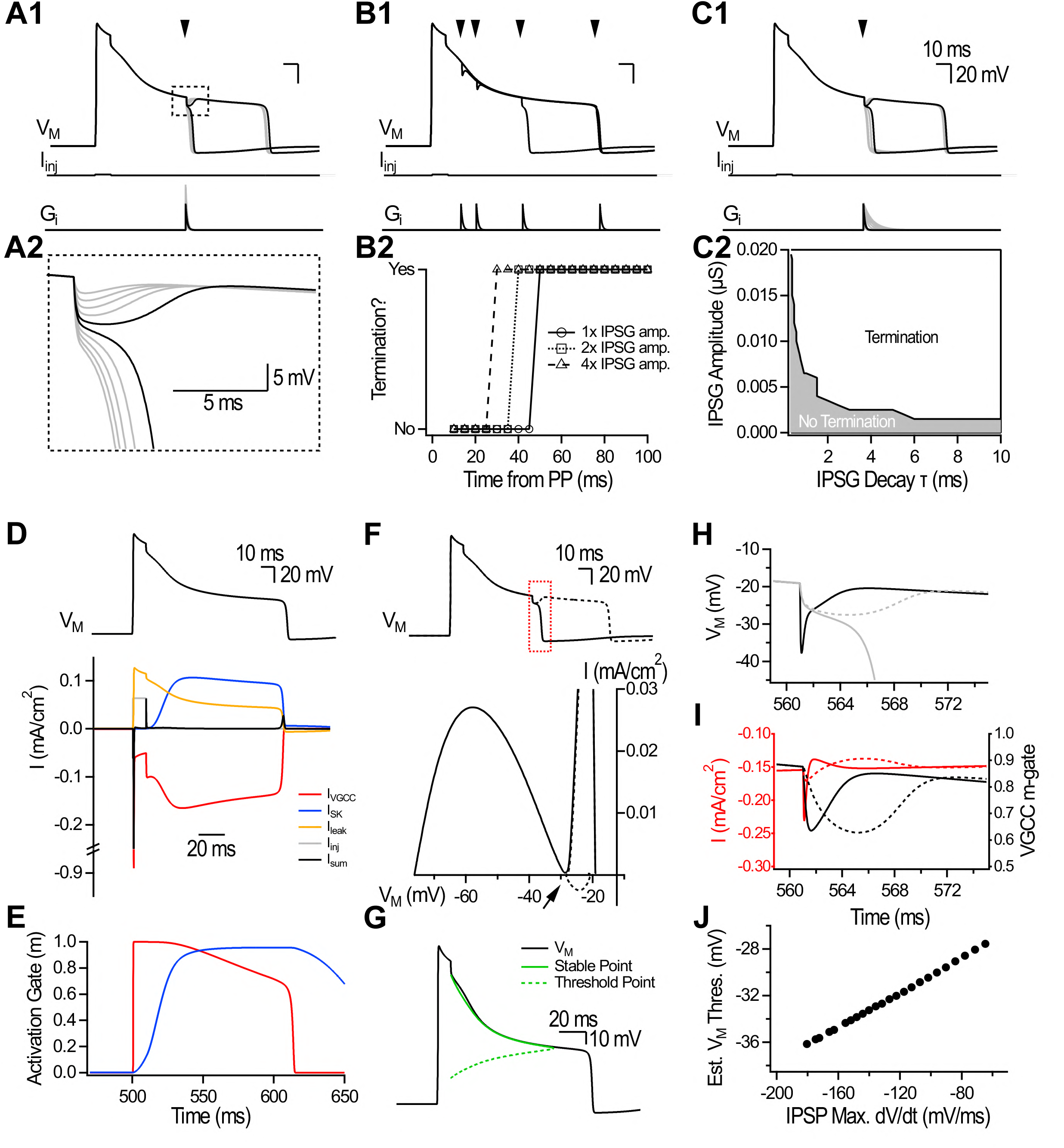
Single-compartment conductance-based model reveals the biophysical mechanisms of plateau termination. **A1**, Simulations showing plateaus generated by brief current injection. An IPSG was delivered 50 ms following plateau initiation. Gradually increasing the IPSG amplitude revealed all-or-none termination. **A2**, Blown-up boxed region from A1 shows the transition from non-terminating to terminating IPSPs. **B1**, Constant IPSG delivered at 10, 20, 50 and 100 ms (indicated by arrowheads) during the plateau. The IPSG amplitude was initially adjusted to the just-threshold value for plateau termination at 50 ms. **B2**, Termination as a function of IPSG timing (5 ms increment) for the data presented in B1, and for IPSGs with 2x (squares) and 4x (triangles) amplitude. **C1**, Plateaus were challenged with IPSG of increasing decay τ. **C2**, Results from simulations in which the parameter space for IPSG amplitude and decay τ was explored. **D**, Dissection of currents (bottom) during simulated plateau (top). Currents are color-coded according to the legend in the figure. **E**, Activation gate (m) for the VGCC (red) and SK (blue) conductances. **F**, Just-subthreshold (dotted line) and just suprathreshold (full line) IPSPs evoked by an IPSG delivered at 50 ms during the plateau. I-V curves for the two traces shown above, focusing on the region identified by the red box starting at the IPSP initiation. The arrow points to the x-axis crossing, a repelling point in V_M_. **G**, Solving the HH equation for I = 0 reveals two sets of V_M_ values within the plateau physiological range. Inferred stable point (green full line) and inferred threshold point (green dotted line). **H**, Simulations specifically selected to show how a fast IPSP (black line) can reach V_M_ values much more hyperpolarized without terminating the plateau than a slower IPSP. For the slow IPSP, the just subthreshold (dotted gray line) and just suprathreshold (full gray line) conditions are shown. **I**, VGCC current and m-gate evolution for the non-terminating examples presented above. **J**, Estimated termination threshold as a function of the IPSP maximum dV/dt. This estimation is obtained from pairs of just subthreshold and just suprathreshold values, where the estimated termination threshold corresponds to the most hyperpolarized V_M_ value reached during the just subthreshold IPSP, and the IPSP maximum dV/dt is measured from the just suprathreshold IPSP.

First, an inhibitory conductance (IPSG) was injected at 50 ms during the plateau (Fig. 6A1). Gradually increasing IPSG amplitude revealed all-or-none termination. Failures to terminate plateaus were characterized by an IPSP waveform and plateaus continued their course seemingly unaffected (Fig. 6A1 – A2). Second, the IPSG amplitude was adjusted at the just-threshold value to terminate plateaus at 50 ms and then injected at multiple timepoints in 5 ms increments from plateau generation (Fig. 6B1 – B2). Plateaus were initially resistant to inhibition but became increasingly susceptible to termination as they progressed. IPSGs with larger amplitudes terminated plateaus earlier (Fig. 6B2). Third, we systematically swept through pairs of IPSG amplitude and decay time constant (τ) to identify multiple parameter pairs sufficient to terminate plateaus (Fig. 6C1 – C2). For a given IPSG amplitude, slower IPSGs were more likely to terminate plateaus (Fig. 6C2). Therefore, a simple model incorporating VGCC and SK conductances recapitulated the main experimental findings.

We next used the model to understand plateaus in the absence of inhibitory events. Dissection of current dynamics revealed that during the plateau, I_VGCC_ was pitted against I_SK_ and I_leak_, such that the outward and inward currents nearly cancelled each other (Fig. 6D). I_VGCC_ sustained V_M_ depolarization, which in turn kept VGCC activated, forming a positive feedback loop. As [Ca^2+^]_i_ rose, I_SK_ gradually increased, V_M_ slowly hyperpolarized and I_sum_ trended closer to the x-axis without crossing it. This interaction suggested a near-stable point in V_M_. Plateaus ended spontaneously when V_M_ exited the range of value allowing sustenance of the positive feedback loop. The end of the plateau was characterized by a sharp drop in the VGCC m-gate and a large outward current unmasked by the abrupt loss of I_VGCC_ (Fig. 6E). These results indicate that plateaus are a near-stable point in V_M_ and that VGCC deactivation marks the end of the plateau.

We analyzed inhibition-driven terminations by comparing supra- and subthreshold IPSPs (Fig. 6F). In simulations with suprathreshold IPSPs, the I-V trajectories approached but did not cross the x-axis, eventually producing a net outward current that brought V_M_ back to its resting value. Like spontaneously ending plateaus, this was characterized by a sharp drop in the VGCC m-gate and unmasking of a large outward current (Fig. 6F and Fig. S5A – C). In contrast, subthreshold IPSPs showed I-V trajectories that crossed the x-axis, reversed slope and returned V_M_ to the plateau state. This suggested the existence of a repelling (threshold) point in V_M_ that underlies all-or-none terminations (Fig. 6F). The reversal described above occurred because the increased Ca^2+^ driving force and reduced K^+^ driving force rebalanced the currents, opposing the IPSC and restoring the plateau (Fig. S5A – F). When the IPSC did not sufficiently disrupt this balance, m could catch up to its steady-state value (m_inf_), sustaining the plateau (Fig. S5F). Thus, for IPSCs to terminate plateaus, they must overcome this dynamic rebalancing to break the positive feedback loop maintaining VGCC activation. These findings provide a mechanistic explanation for all-or-none plateau termination.

Co-existence of stable and threshold points describes plateau behavior in cardiac Purkinje fibers (54). To examine their evolution during plateaus in CA1-PYRs, we solved the Hodgkin-Huxley equation for I = 0 by assuming steady-state conditions (Fig. 6G). We identified two sets of gradually evolving V_M_ values within the plateau range that balanced the total membrane current (Fig. 6G). The first set of values (solid green line) was tracked by V_M_, consistent with the idea that plateaus are a gradually hyperpolarizing near-stable point in V_M_. The second set (dotted green line) initially resided at more hyperpolarized levels but gradually drifted towards more depolarized levels. This is the approximative threshold point, and its evolution explains why plateaus display an early resistance to inhibition that gradually fades.

Why are slow IPSPs better at terminating plateaus than fast IPSPs (Fig. 6H)? We observed that fast IPSPs can reach a much more hyperpolarized V_M_ without terminating plateaus than slow IPSPs (for which the just-subthreshold trace is shown, dotted line, Fig. 6H). As described above, IPSPs hyperpolarize V_M_ which rebalances the currents. Because VGCC m-gate updating lags (and particularly so in that V_M_ range, Fig. S5G), m remains high despite the hyperpolarized V_M_, and I_VGCC_ increases (Fig. 6I). This rebalancing adds to the inward current that the IPSC must overcome to terminate the plateau. Slow IPSPs do not experience this added current load, because m evolves with the IPSP (Fig. 6I). This interaction generates a dynamic threshold for plateau termination that collapses when approached with a fast V_M_ trajectory (Fig. 6J). Overall, these results provide a mechanistic explanation for why slow and sustained inhibition, such as that provided by OLM^Ndnf^, favors plateau termination.

### Plateau termination transforms binary events into graded dendritic Ca^2+^ signals

Large dendritic Ca^2+^ influxes are associated with plateaus, including events supporting BTSP (4, 40, 55, 56). We next evaluated the impact of plateau termination on dendritic Ca^2+^ elevations.

We used random-access two-photon Ca^2+^ imaging to visualize Ca^2+^ elevations in CA1-PYR dendritic arbors. CA1-PYRs were filled with a morphological indicator (Alexa-594, 40 µM) and the medium-affinity Ca^2+^ indicator Fluo-5F (385 µM) (Figs. 7A – C and S6; and see Materials and Methods for description of the recording sites). Plateaus generated Ca^2+^ transients (CaTs) that were larger and lengthier than CaTs evoked by backpropagating APs (plateau: 4 ± 0.4 ΔF/F; APs: 0.9 ± 0.1 ΔF/F; n = 57 dendritic sites across N = 4 neurons, p < 0.001, Wilcoxon signed-rank test; Fig. 7C – D), yielding a significantly larger area under the curve (AUC) for plateau-evoked CaTs (plateau: 0.57 ± 0.06 ΔF/F*s; APs: 0.06 ± 0.01 ΔF/F*s; n = 57 dendritic sites across N = 4 neurons, p < 0.001, Wilcoxon signed-rank test; Fig. 7E). Plateaus were isolated from APs with TTX, in which case plateaus still evoked large CaTs across the dendritic arbor (Fig. 7F). The duration of the Ca^2+^ transients generally mirrored that of the plateaus recorded electrophysiologically, consistent with maintenance of VGCCs in the activated state (Fig. 7F). Plateau-evoked CaTs were observed at long-distances from the soma in primary and secondary branches in the presence of TTX (n = 106 dendritic sites across N = 6 neurons; Fig. 7G).

**Figure 7.**
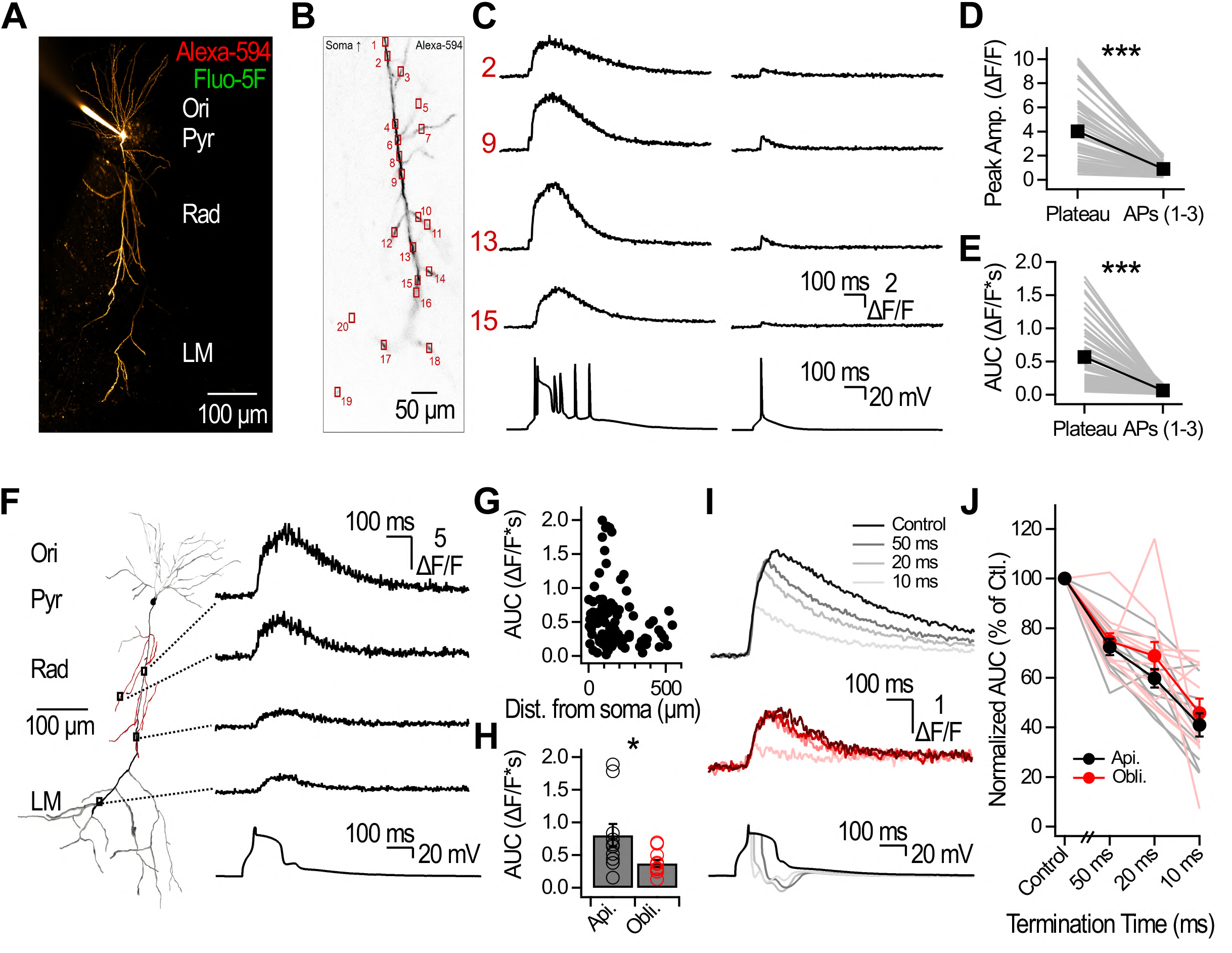
Plateau termination shapes dendritic Ca^2+^ transients. **A**, Montage from maximal projections of two-photon Z-stacks of a CA1-PYR filled with Alexa-594. **B**, All recording sites selected on the dendritic arbore for the exemplars presented in C. **C**, Dendritic Ca^2+^ transients evoked by AP and plateaus or by single backpropagating AP. The imaging speed was 555 Hz. Numbers correspond to the physical positions shown in B, and all recording sites for that example are shown in Fig. S6. Ca^2+^ transients amplitude (**D**) and AUC (**E**) evoked by APs and plateaus or by 1 – 3 APs without plateaus. Individual lines correspond to a dendritic recording site. Average ± SEM is shown in black. **F**, Plateau-associated Ca^2+^ transients evoked in the presence of TTX are observed throughout the dendritic arbor. **G**, Plateau-evoked Ca^2+^ transients recorded in presence of TTX as a function of distance from the soma. **H**, Plateau-evoked Ca^2+^ transients in the presence of TTX measured in apical and in obliques dendrites. **I**, Examples showing Ca^2+^ transients recorded in apical (black) and in oblique (red) dendrites associated with control plateaus, and at indicated termination times. **J**, Normalized Ca^2+^ transient AUC as a function of plateau termination time. Data was normalized to the control plateau.

We focused the analysis on apical and oblique dendrites within stratum radiatum, which are the primary target sites of Schaffer collateral inputs. Plateau-evoked CaTs were larger in apical dendrites than in the obliques (apical: 0.8 ± 0.17 ΔF/F*s, n = 10 segments; oblique: 0.37 ± 0.05 ΔF/F*s, n = 11 segments; p < 0.05; Mann Whitney U test, both recorded across N = 6 neurons). Lastly, we probed the impact of plateau termination on dendritic CaTs (Fig. 7I).

Plateaus were terminated with mock IPSPs at 10, 20, and 50 ms, and the AUC of plateau-evoked CaTs was compared to control non-terminated plateaus in paired measurements (Fig. 7I – J). Plateau termination diminished CaTs relative to control such that shorter plateaus gave rise to smaller CaTs in apical and oblique dendrites (apical: n = 10 segments; oblique: n = 11 segments, both recorded across N = 6 neurons; Fig. 7J). Therefore, these results show that plateau termination effectively grades dendritic CaTs.

## Discussion

Our results show that plateaus recorded in CA1-PYRs are terminated in an all-or-none manner by dendritic inhibition. Plateaus are initiated in a binary manner, but unlike action potentials, they are sensitive to ongoing synaptic activity. Plateau duration is thus a function of CA1-PYRs intrinsic properties and feedback inhibitory circuit connectivity motifs. Dendritic Ca^2+^ imaging showed that plateau termination controls this critical signal for many forms of synaptic plasticity directly at sites where many excitatory synapses reside. Given that both plateaus and the feedback inhibitory circuit are involved in synaptic plasticity and memory (5, 28, 31, 34, 57–60), we first consider the consequences of plateau termination within a broader circuit context and branch-specific dendritic processing.

All-or-none plateau termination by feedback inhibition may promote sparse ensemble formation by limiting Ca^2+^-dependent plasticity in select neurons, even among those sharing common excitatory drive. The local inhibitory circuit organizes CA1-PYRs into functional ensembles for spatial representation, and feedback inhibition constrains ensemble size (61–64). OLM^α2^ firing is reduced in novel environments, likely promoting the assignment of place cell identity (34). Global suppression of feedback inhibition enhances local population spiking and increases the amplitude and duration of ripples (63). Sharp-wave ripples (SPW-Rs) are high-frequency oscillations that are important for spatial memory and involve coordinated neuronal firing and inhibitory interactions (65–67). The co-occurrence of complex Ca^2+^ spikes in CA1-PYR dendrites with SPW-Rs supports the idea that independent representations may be achieved through the combination of neuron-specific plateau termination thresholds and inhibitory connectivity motifs (16, 68, 69).

We further hypothesize that local plateau terminations spatially confine Ca^2+^ elevations to constrain synaptic plasticity in select branches in an input-dependent manner (15, 28, 34). Individual CA1-PYRs can support multiple spatial representations, likely through dendritic compartmentalization that enables branch-specific encoding (40, 70–72). Plateau compartmentalization allows individual neurons to detect input sequences over behaviorally relevant timescales (73). Plateaus evoked and recorded in single CA1-PYR dendritic branches can exceed 100 milliseconds and spread to neighboring branches (3, 4). Correlated dendritic plateaus in response to heterogeneously tuned inputs may initially support branch-specific encoding, with inhibition possibly guiding local plastic changes to establish sparse and selective representations within a CA1-PYR neuron (64).

### Reconstruction of plateau termination

This study adds to the growing body of evidence that dendritic spikes and plateaus are highly sensitive to inhibition and membrane hyperpolarization (6, 7, 9, 13–15). In CA1-PYRs, our electrophysiological recordings, Ca^2+^ imaging, and computational modeling collectively demonstrate that plateaus arise from VGCC activation to generate a Ca^2+^ current that maintains V_M_ depolarization through a positive feedback loop. Ca^2+^ activates I_SK_, forming a system where two currents of opposing polarity nearly balance each other, consistent with previous pharmacological dissection of plateaus in CA1-PYRs (3, 4). The small net outward current slowly repolarizes V_M_ until a threshold for rapid repolarization is reached, given by the point where the inward current can no longer sustain the positive feedback loop, and VGCC deactivation abruptly repolarizes V_M_ back to the resting state. Synaptic inhibition adds to the outward current pressure and forces plateau termination. This addition is complicated by the IPSP-driven rebalancing of intrinsic conductances, which generates a dynamic threshold point.

Gradually evolving stable and threshold points explain all-or-none plateau repolarization in CA1-PYRs, in a similar manner to cardiac Purkinje fibers and neocortical pyramidal cells (54, 74, 75). Our results further show that the dynamic evolution of the threshold on two timescales governs the impact of synaptic inhibition in CA1-PYRs. The threshold approximated under steady-state conditions shifts towards a more depolarized value during the plateau, conferring an initial resistance to inhibition that gradually dissipates. For a subthreshold IPSP, V_M_ hyperpolarization increases the driving force for Ca^2+^ and increases the current which repolarizes V_M_ back to the plateau phase. This is because VGCC m gate updating is particularly slow in that V_M_ range and lags rapid hyperpolarization (Fig. S5G). The interaction generates a dynamic threshold that collapses when approached with an elevated dV/dt. In a way, this is the inverse of action potential generation, where large but slow depolarization may not trigger firing due to neuronal accommodation (76). Termination threshold is not a fixed V_M_, but rather a dynamically evolving point governed by prior activity and intracellular Ca^2+^ dynamics. This explains why slow IPSPs mediated by OLM^Ndnf^ more effectively terminate plateaus than the faster IPSPs generated by OLM^α2^. It is anticipated that plateau termination by synaptic inhibition is broadly applicable across systems and species, given the prevalence of VGCCs in mediating multiple forms of dendritic plateaus (75, 77).

### Intrinsic mechanisms supporting plateaus

Plateaus in this study were of long duration, but well within the range of plateaus recorded *in vivo* with K^+^ internal solution (5, 40, 41). Plateaus in acute slices are made evident by using Cs^+^-based intracellular solution (5), cholinergic modulation (17), coincident excitatory inputs activation (1), dendritic glutamate uncaging (3, 4, 78), and high activity levels (79). While excitatory afferents acting at multiple synapses and receptors can provide the depolarization required for plateau initiation and are essential in understanding the intricate interactions supporting synaptic plasticity, this aspect is not the focus here (1, 5, 80, 81). We described how intrinsic conductances suffice to generate and sustain plateaus in CA1-PYRs, as revealed across the dendritic arbor using random-access two-photon Ca^2+^ imaging.

Our relatively simple single-compartment model captures a phenomenon acting across the complex CA1-PYR dendritic arbor that contains multiple types of heterogeneously distributed channels (82–84). While the model explains the main determinants of plateau termination, differences between experiments and model remain. We and others previously observed that plateaus end spontaneously in apamin, are shortened by TTX, can be regenerated and ride on a depolarizing envelope. Our results further show that IPSPs become larger when delivered later during plateaus despite constant optogenetic stimulation, a feature not captured by the model. Additional mechanisms and conductances not accounted for in our model likely contribute to support these features (4, 17, 85–91). Our latter observation requires closer attention, as it could enhance the plateaus’ early resistance to synaptic inhibition. A Ca^2+^-activated Cl^-^ current contributes to action potential repolarization in CA1-PYRs and is likely to be amplified by the larger Ca^2+^ elevations generated by plateaus measured here (92). Decaying intracellular Cl^-^ concentration during the plateau provides an attractive explanation for the growing IPSP amplitude not associated with changes in input resistance. Depending on the distribution of this current across the dendritic arbor, it may confer resistance to inhibition in select compartments or even contribute to early plateau termination altogether.

### Structure, function, and timing of specialized feedback inhibitory elements

The timing of inhibition determines whether plateaus are terminated or allowed to persist, which focuses our attention on the recruitment of feedback inhibitory neurons. Repetitive CA1-PYR spiking optimally recruit interneurons including OLM^Ndnf^ and OLM^α2^ where short-term facilitation allows for a single CA1-PYR to drive feedback interneurons (22, 23, 35, 42, 93). Delayed post-integration recruitment is aligned with the role of *Sst*-INs in regulating CA1-PYR activity toward the end of place fields or during burst firing (30). Plateaus themselves can provide the signal for a CA1-PYR to recruit feedback inhibition by triggering axonal spikes and glutamate release from CA1-PYRs despite absence of measurable somatic firing (94). Furthermore, multiple CA1-PYRs form synapses onto a single feedback interneuron. OLMs receive over 5,000 excitatory synapses, mostly from local CA1 collaterals which usually form a single contact site per cell (19, 95–97). Therefore, multiple modes of activity in the CA1 circuit are expected to recruit feedback interneurons, crucial for terminating plateaus.

Our experiments and modeling results converge to show why slower inhibitory currents drive plateau termination more effectively, providing a distinct advantage to OLM^Ndnf^ over OLM^α2^ for terminating plateaus with more efficacy or at earlier timepoints. The importance of OLM^Ndnf^ for plateau termination and inhibitory control of CA1-PYRs is further amplified by our recent demonstration that OLM^Ndnf^ but not OLM^α2^ preferentially target CA1-PYRs (35). While OLM^Ndnf^ and OLM^α2^ are transcriptomically similar and exhibit a common anatomical phenotype (35, 36), our data demonstrated only minimal overlap between the two subpopulations in the transgenic animals used. Anatomical analysis confirmed our previous observations that OLM^Ndnf^ axons penetrate LM less than those of OLM^α2^, arguing that dendritic filtering is unlikely to account for the differences in current kinetics reported here (35). Slower IPSCs mediated by OLM^Ndnf^ may originate from different release machinery and/or postsynaptic receptor expression. This observation, combined with the fact that plateau termination is a cell-intrinsic mechanism, supports the general principle that any sufficiently strong and slow inhibition can trigger terminations to control dendritic Ca^2+^ elevations.

## Materials and Methods

### Animals

All experiments on animals were approved by IACUC at the University of Pittsburgh. Animals were given access to food and water ad libidum. Experiments were performed on animals of either sex. Animals used for experiments were between P25 and P65. The following transgenic animals were used in this study: *Sst-Cre;;Ai32*, *Chrna2-Cre;;Ai9*, *Ndnf-FlpO;;Nkx2-1-Cre;;Ai80*, *Chrna2-Cre;;Sst-FlpO;;Ai80*, *Ndnf-FlpO;;Nkx2-1-Cre;;Ai65*, *Ndnf-FlpO;;Chrna2-Cre;;Ai224*.

Littermates from these crosses were also used in experiments. The animals were obtained by breeding the following animals: *Sst-Cre* (Sst^tm2.1(cre)Zjh^/J; JAX: 013044) were maintained as homozygous (98); *Sst-FlpO* (B6J.Cg-Sst^tm3.1(flpo)Zjh^/AreckJ; JAX: 031629) were maintained as homozygous (99); *Chrna2-Cre* were maintained as hemizygous (gift from Dr. Klas Kullander) (28). *Ndnf-FlpO* were maintained as heterozygous (Dr. Robert Machold, NYU) (35). *Nkx2-1-Cre* (C57BL/6J-Tg(Nkx2-1-cre)2Sand/J; JAX: 008661) were maintained as hemizygous (100). *Ndnf-FlpO;;Nkx2-1-Cre* were maintained as het/hemi. *Chrna2-Cre;;Sst-FlpO* were maintained as hemi/hom. The following reporter lines were used and maintained as homozygous:

Ai9 (B6.Cg-Gt(ROSA)26Sor^tm9(CAG-tdTomato)Hze^/J; JAX: 007909) (101).

Ai32 (B6.Cg-Gt(ROSA)26Sor^tm32(CAG-COP4*H134R/EYFP)Hze^/J; JAX: 024109) (102).

Ai65 (B6;129S-Gt(ROSA)26Sor^tm65.1(CAG-tdTomato)Hze^/J; JAX: 021875) (103).

Ai80 (B6.Cg-Gt(ROSA)26Sor^tm80.1(CAG-COP4*L132C/EYFP)Hze^/J; JAX: 025109) (104).

Ai224 (B6.Cg-Igs7^tm224(CAG-EGFP,CAG-dTomato)Tasic^/J; JAX: 037382) (50).

### Acute hippocampal slice preparation

Animals were deeply anesthetized with isoflurane using the drop-jar method before decapitation. The brain was rapidly extracted and immersed in oxygenated (95% O_2_, 5% CO_2_) ice-cold sucrose-based artificial cerebrospinal fluid (sucrose aCSF) containing (in mM): 185 sucrose, 25 NaHCO_3_, 2.5 KCl, 25 glucose, 1.25 NaH_2_PO_4_, 10 MgCl_2_, and 0.5 CaCl_2_; pH 7.4, 300 mOsm. Brain hemispheres were glued on a specimen disk. Acute slices (300 µm) were prepared on a vibratome (VT1200 S, Leica) and the slicing chamber was maintained on ice and continuously oxygenated. Slices were allowed to recover in oxygenated and heated (32°C) sucrose aCSF for 30 minutes, before being transferred to oxygenated and heated (32°C) recording aCSF that contained (in mM): 125 NaCl, 25 NaHCO_3_, 2.5 KCl, 10 glucose, 2 CaCl_2_, and 2 MgCl_2_; pH 7.4, 300 mOsm. The bath temperature was turned off upon transfer to recording aCSF, and the solution was left to cool down to room temperature. Slices were allowed to recover for 30 minutes before experiments began, and acute slices were maintained at room temperature and continuously oxygenated for up to 7 hours following the slicing procedure.

### Electrophysiology

Acute slices were transferred to a recording chamber, held under a harp, and continuously perfused with warm (32°C) and oxygenated recording aCSF. The CA1 region of the hippocampus was identified under an upright microscope (Scientifica) with a 4X objective (NA = 0.1, Olympus) and neurons were identified under a 40X water-immersion objective (NA = 0.8, Olympus) for visually guided whole-cell recordings. Recording electrodes were fabricated from borosilicate glass capillaries (BF150-86-10HP, Sutter Instrument) on a P-1000 micropipette puller (Sutter Instrument). Recording electrodes were filled with a solution containing: 130 Cs-methanesulfonate, 10 HEPES, 2 MgCl_2_.6H_2_O, 4 Mg_2_ATP, 0.3 NaGTP, 7 Phosphocreatine di(tris), 0.6 EGTA, and 5 KCl; pH 7.3 and 290 mOsm. Electrodes filled with this solution had a resistance of 4 – 8 MΩ when inserted in the bath. The liquid junction potential was not corrected. Micromanipulators (Patchstar, Scientifica) were used to approach the recording electrodes to targeted neurons and standard techniques were used to obtain whole cell recordings. The electrophysiological signal was amplified (Multiclamp 700B, Molecular Devices) and digitized at 10 kHz (Digidata 1550B, Molecular Devices) before displaying and saving to a personal computer (Clampex v11.3, Molecular Devices). Only neurons for which the access resistance was < 30 MΩ and with a resting membrane potential more hyperpolarized than −55 mV were included. Optogenetic stimulation was delivered from a pE-400 max CoolLED illumination system at 450 nm (CoolLED Ltd.) Light power was measured under the objective with a PM101R power meter (Thor Labs). Salts used to prepare solutions for electrophysiological experiments were purchased from Sigma except for NaHCO_3_ (Fisher Scientific). The following compounds were also used in experiments: TTX (1 µM, tetrodotoxin with citrate, Biotium), apamin (100 nM, Medchemexpress LLC) and CdCl_2_ (100 µM, Thermo Scientific Chemicals).

### Two-photon Ca^2+^ imaging

Illumination was provided by a Chameleon Discovery NX femtosecond laser (80 MHz repetition rate, 100 fs pulse duration, Coherent) tuned at 810 nm. The laser was directed to a random-access two-photon microscope (AODScope, Karthala System). The laser position was controlled by a pair of acousto-optic deflectors in the X-Y planes and focused under a 25X water-immersion objective (numerical aperture = 0.95, Leica). Transmitted and emitted photons were detected below and above the sample, each split in a green and a red channel and collected with four photomultiplier tubes (PMTs). For transmitted and emitted signal, photons were filtered through an IR blocking filter (TF1, Thorlabs) and split into two channels using a 562 nm dichroic mirror (Semrock). Green channel photons were filtered with a 510/84 nm bandpass filter and red channel photons were filtered with a 607/70 nm bandpass filter before being directed to two H12056P-40 PMTs (Hamamatsu) operating in photon counting mode. Transmitted and emitted signals were averaged in their respective channels before analyzing the data. Whole-cell recordings were obtained from CA1-PYRs using the Cs^+^-based intracellular solution detailed above and supplemented with Alexa-594 (40 µM) and Fluo-5F (385 µm). Recording sites were distributed across the dendritic arbor in a field of view. We typically recorded from 20 sites near-simultaneously with a 90 µs dwell time, yielding an average imaging speed of 500 Hz, similar to our previous studies employing random-access two-photon microscopy (105–107). Plateaus were terminated with mock IPSPs. We observed that plateaus were generally harder to terminate in these conditions, possibly because of the added Ca^2+^ buffering capacity preventing I_SK_ activation. We used mock IPSPs with larger amplitude and longer offset to ensure terminations. This was also the case in the presence of TTX. Dendritic segments corresponded to 1 – 4 physical recording sites separated by less than 76 µm across a continuous dendritic branch.

### Single cell labeling, confocal imaging, and anatomical tracing

Neurons filled with biocytin (1 – 3%, w/v) were fixed in 4% PFA dissolved in PBS. Biocytin was revealed in filled neurons using the procedure described before (35, 39). Briefly, slices with biocytin-filled neurons were treated with H_2_O_2_ (30 min), Triton X-100 (1 hour), and a streptavidin-conjugated Alexa-633 (1:100, incubated overnight). Slices were rinsed, mounted on microscope slides with coverslips, and allowed to rest for at least 2 weeks before confocal imaging. Confocal images were obtained on a Leica SP8 or Zeiss Axo Imager.Z2 upright confocal microscope with a 40X oil-immersion objective. Neurons were traced in Neurolucida 360 (v.2024.2.2, MBF Bioscience). The axonal distribution was measured in 10 µm bins starting from the pyramidal cell layer. Only neurons with total axonal length > 450 µm were included for analysis presented in Fig. 4E. For data presented in Fig. 4A – B, images were obtained on a Nikon A1R-HD confocal microscope with a 20x objective at 0.63 pixels/μm in x-y and four μm z-steps. Brains were serially sectioned in the coronal plane, and sections 100 μm apart were analyzed. Data from the left and right hemispheres were combined. Confocal scans encompassed the entirety of the 50 μm thickness of the CA1 hippocampus. Sample preparation, imaging, and cell quantification were performed similarly as described previously (108).

### Data Analysis and statistical treatment

Electrophysiological data was analyzed in Clampfit (v11.3, Molecular Devices), Igor Pro (v9.05, WaveMetrics) and Microsoft Excel (Microsoft). Ca^2+^ imaging data was exported to and analyzed in Microsoft Excel and Igor Pro. Images were analyzed in Fiji (ImageJ 1.54f). Normality of the data distribution was tested using the Shapiro-Wilk test. For normally distributed paired data, Student’s paired t test was used. For non-normally distributed paired data, Wilcoxon signed rank test was used. For normally distributed unpaired data, Student’s unpaired t test was used. For non-normally distributed unpaired data, a Mann Whitney U test was used. Significance levels are reported everywhere as follows: n.s. = non-significant; * p < 0.05; ** p < 0.01; and *** p < 0.001. Statistical tests were performed in Python (Jupyter Notebook, v7.2.2) or in Clampfit.

### Biophysical modelling

A single-compartment conductance-based NEURON (v.8.2.6) model was assembled and controlled in Python (109, 110). The compartment was a sphere with a diameter of 10 µm with surface area of 3.14 x 10^-6^ cm^2^. The membrane capacitance (C_m_) was set to 1 µF/cm^2^. The inhibitory synapse was connected in the middle of the compartment. The following mechanisms were included: passive leak (pas) (109), voltage-gated calcium channel (cal, referred to as VGCC) (82, 83), calcium-activated potassium current (kca) (111), intracellular calcium dynamics (cad) (112), inhibitory synapse (ExpSyn, referred to as GABA) (109). Simulations were performed at 32°C with a time step of 0.025 ms. The differential equation governing V_M_ is given by:

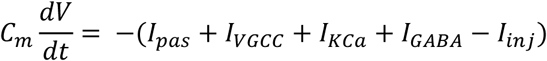

with:

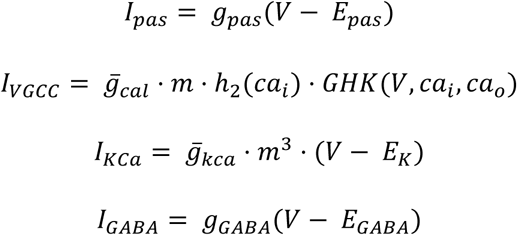

For *I*_*VGCC*_, the differential equations for the gating variables *m*_*cal*_ is:

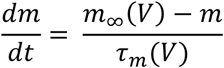

with:

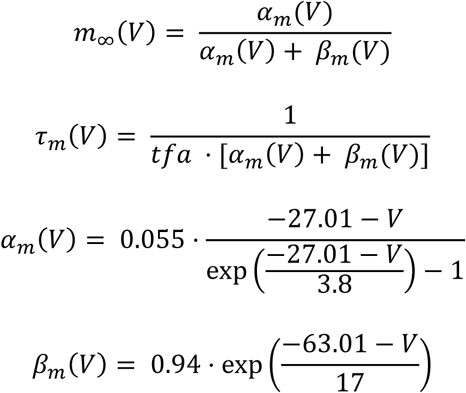

For *I*_*VGCC*_, the Ca^2+^-dependent inactivation factor is given by:

For *I*_*VGCC*_, *GHK*(*V*, *Ca*_*i*_, *Ca*_*o*_) is given by:

For *I*_*KCa*_, the differential equation for the gating variable *m*_*KCa*_ is:

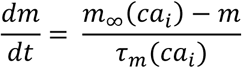

with:

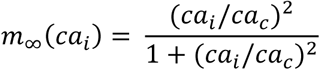

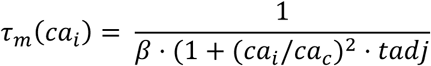

The [Ca^2+^]_i_ dynamics are governed by the differential equation:

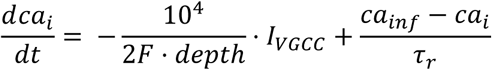

with:

*Ca*_*i*_= [Ca^2+^]_i_

*F* = Faraday constant

*depth* = depth of the submembrane shell in which Ca²⁺ accumulates (μm), here set to 0.1

*Ca*_*inf*_= steady-state [Ca^2+^]_i_

τ_*r*_= Ca^2+^ removal time constant

Conductances for VGCCs and SK were determined empirically. A grid search was performed to find combinations of parameters yielding plateaus with similar duration as observed experimentally (Fig. S3), and these parameters were generally comparable with previous reports (83, 113, 114). The biophysical model will be made available on modeldb at the time of publication.

## Acknowledgements

We thank Dr. Jean-Claude Lacaille for helpful comments and discussion. This work was supported by a R00 Pathway to Independence Award from NIMH (R00MH126157) to SC. RM was supported by NINDS grant P01NS074972. We thank Dr. Klas Kullander for the Chrna2-Cre mouse.

---

**Supplementary Figure 1.**
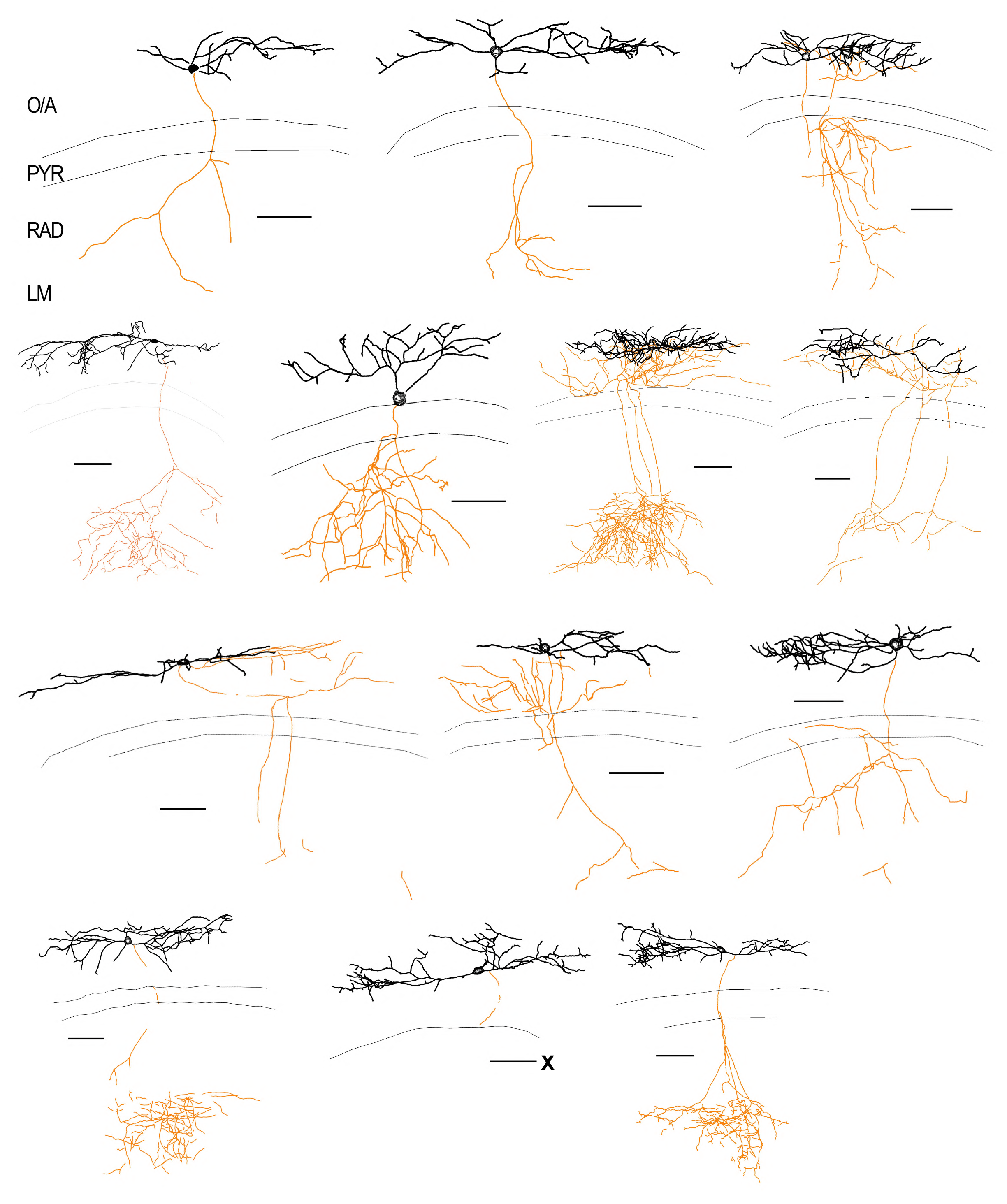
Neurolucida reconstructions of OLM^Ndnf^ recorded in *Ndnf-FlpO;;Nkx2-1-Cre;;Ai65* triple transgenic mice. Soma and dendrites are shown in black, and the axon is shown in orange. Neuron excluded from Figure 4D analysis (total axonal length < 450 µm) is marked with an X next to scale bar. Scale bars = 100 µm.

**Supplementary Figure 2.**
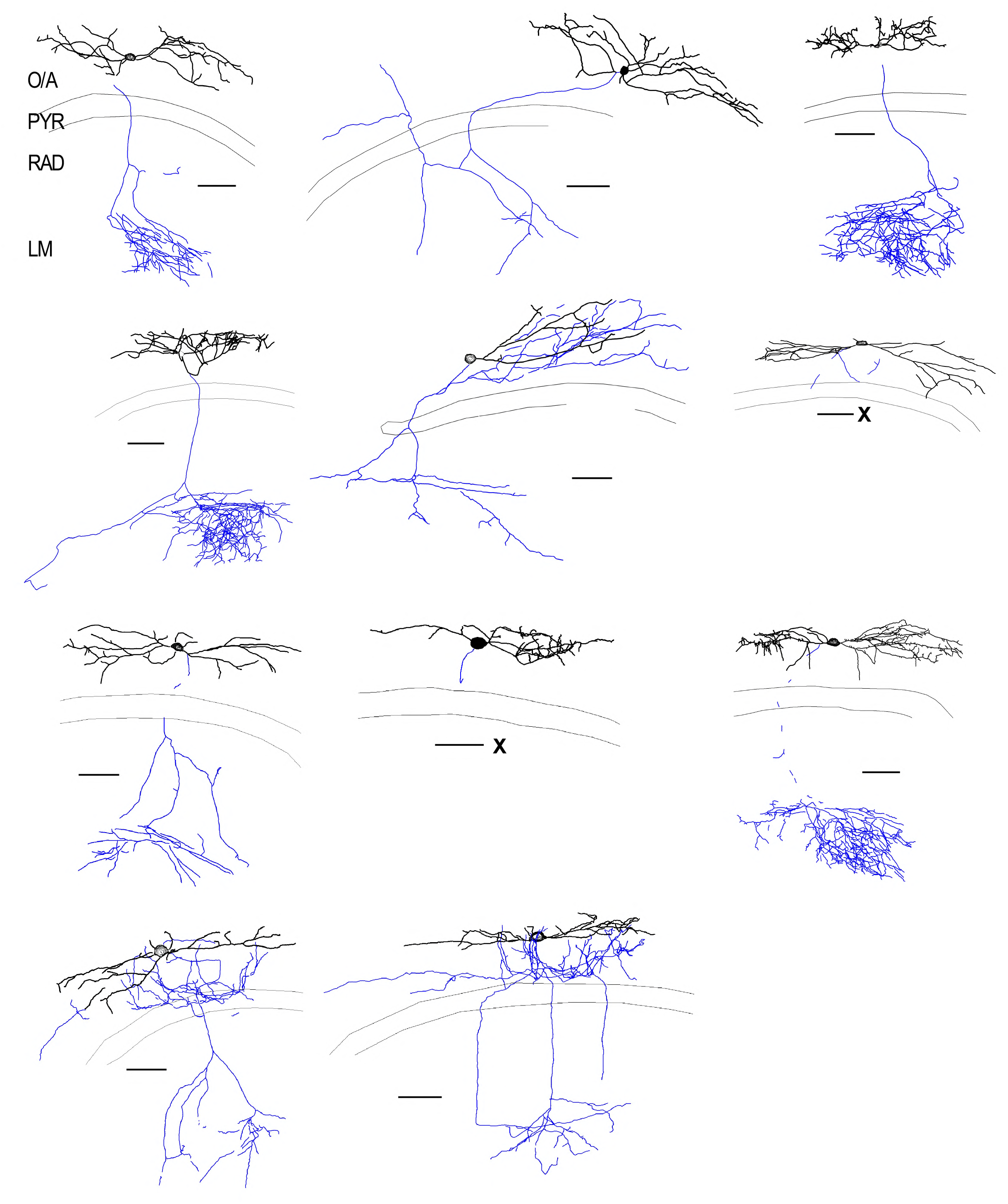
Neurolucida reconstructions of OLM^α2^ recorded in *Chrna2-Cre;;Ai9* double transgenic mice. Soma and dendrites are shown in black, and the axon is shown in blue. Neurons excluded from Figure 4D analysis (total axonal length < 450 µm) are indicated with an X next to the scale bar. Scale bars = 100 µm.

**Supplementary Figure 3.**
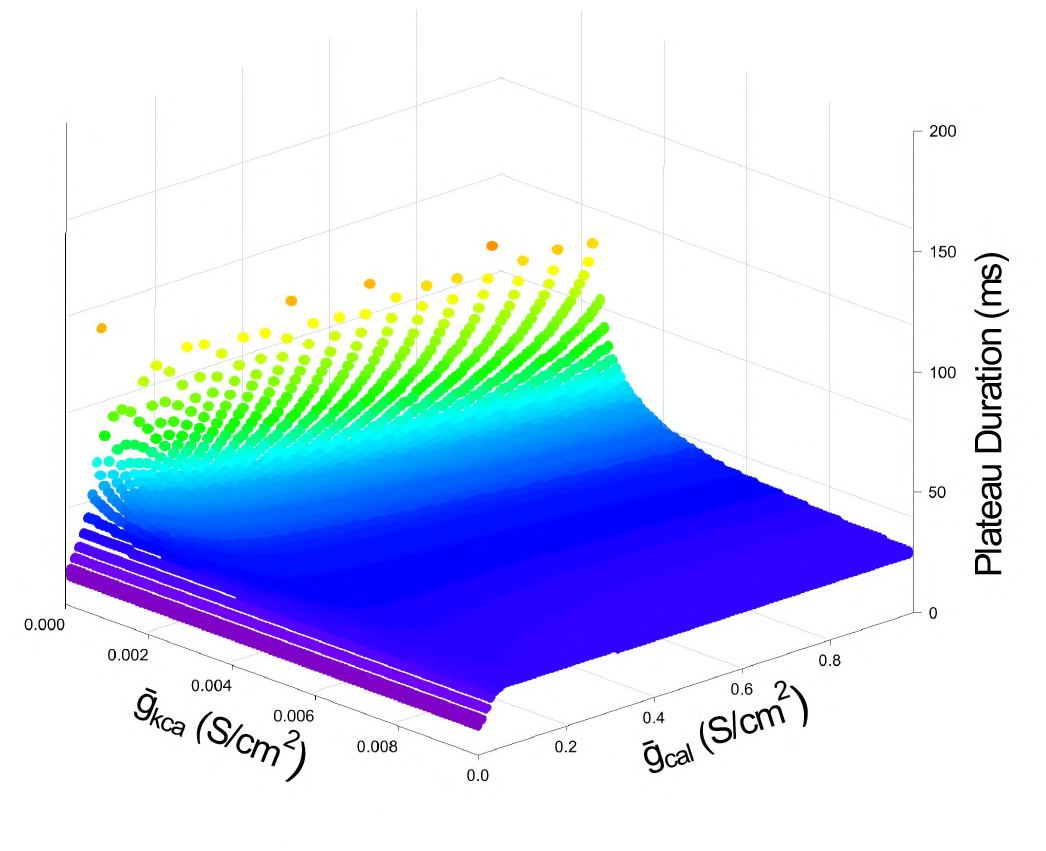
Plateau duration as a function of VGCC and SK conductances in the model. Warmer colors code for longer plateaus.

**Supplementary Figure 4.**
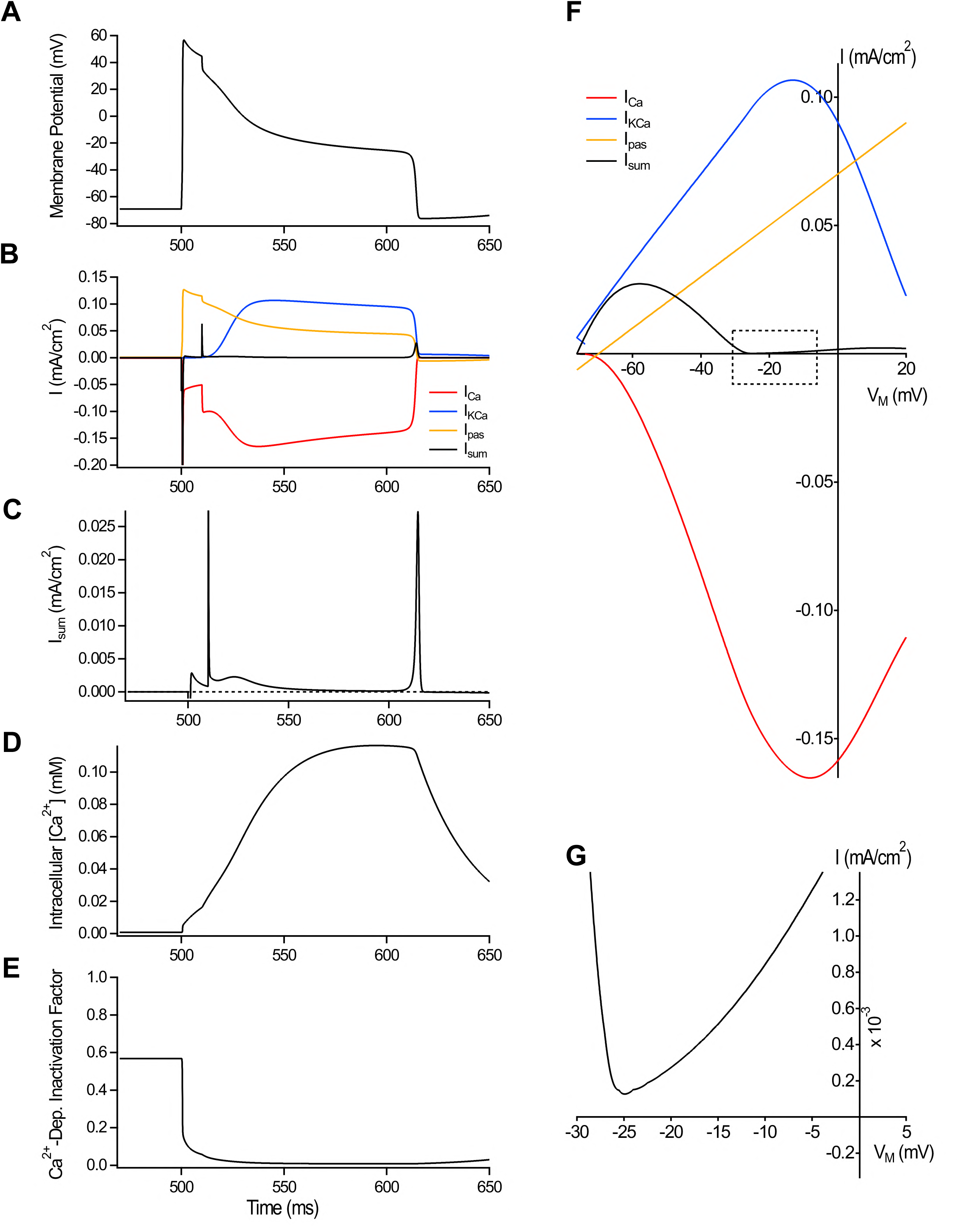
Evolution of currents, [Ca^2+^]_i_ and Ca^2+^-dependent inactivation factor during plateaus. **A**, Simulated plateau with associated individual currents (**B**), total current (**C**), [Ca^2+^]_i_, (**D**) and Ca^2+^-dependent inactivation factor for the VGCC (**E**). Examples correspond to that of Fig. 6D – E. **F**, I-V profile of currents during plateaus. **G**, I-V profile of the sum of current, zoomed in from the boxed region in F to show that the I-V trajectory approaches but doesn’t cross the x-axis for spontaneous plateau termination.

**Supplementary Figure 5.**
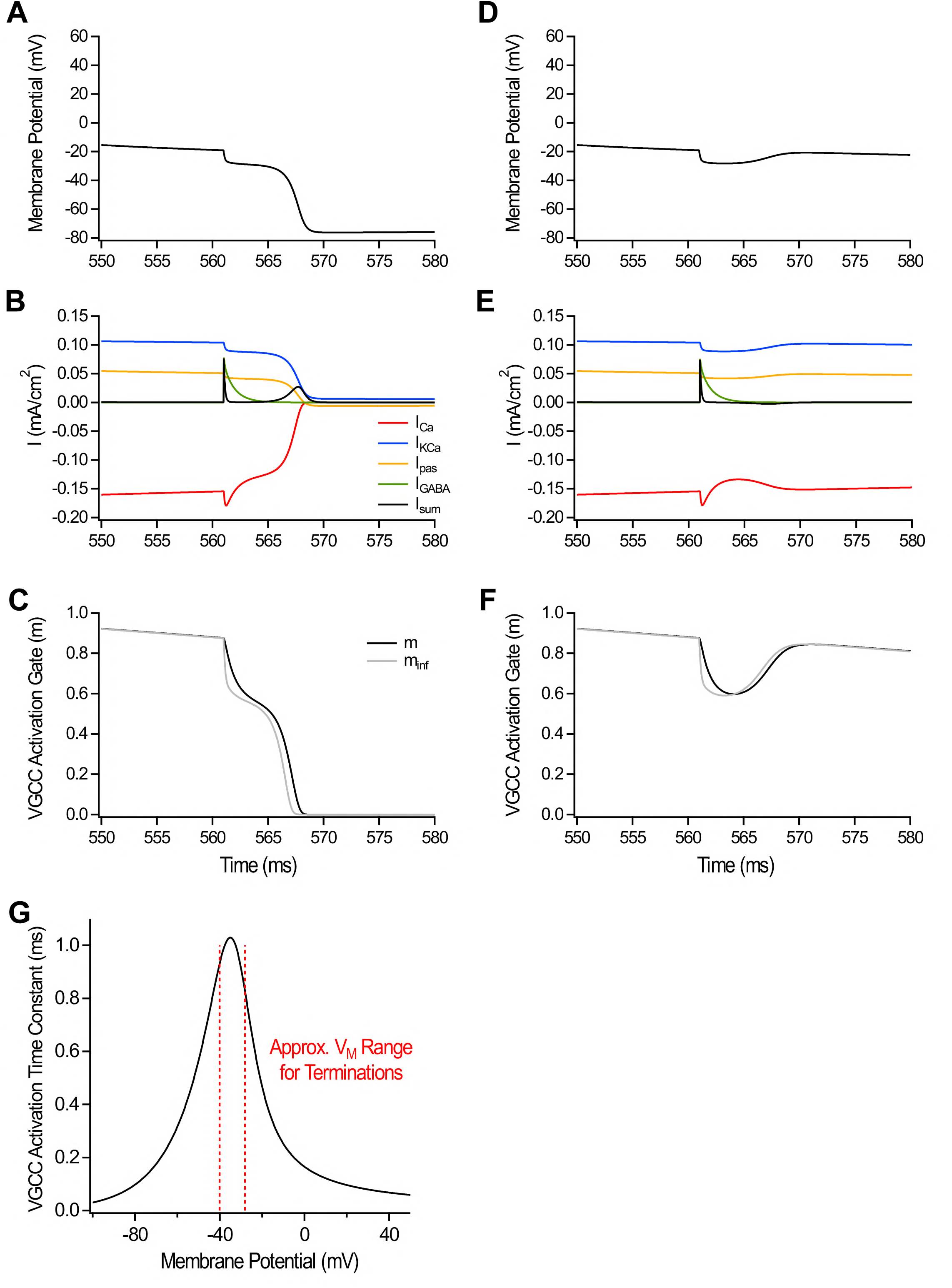
Evolution of currents and VGCC m-gate dynamics during plateaus for subthreshold and suprathreshold IPSPs. **A – C**, Membrane potential (A), individual currents (B) and VGCC m-gate (C) as a function of time for supra-threshold IPSP. **D – F**, Membrane potential (D), individual currents (B) and VGCC m-gate as a function of time for subthreshold IPSP. **G**, VGCC m-gate activation time constant as a function of membrane potential.

**Supplementary Figure 6.**
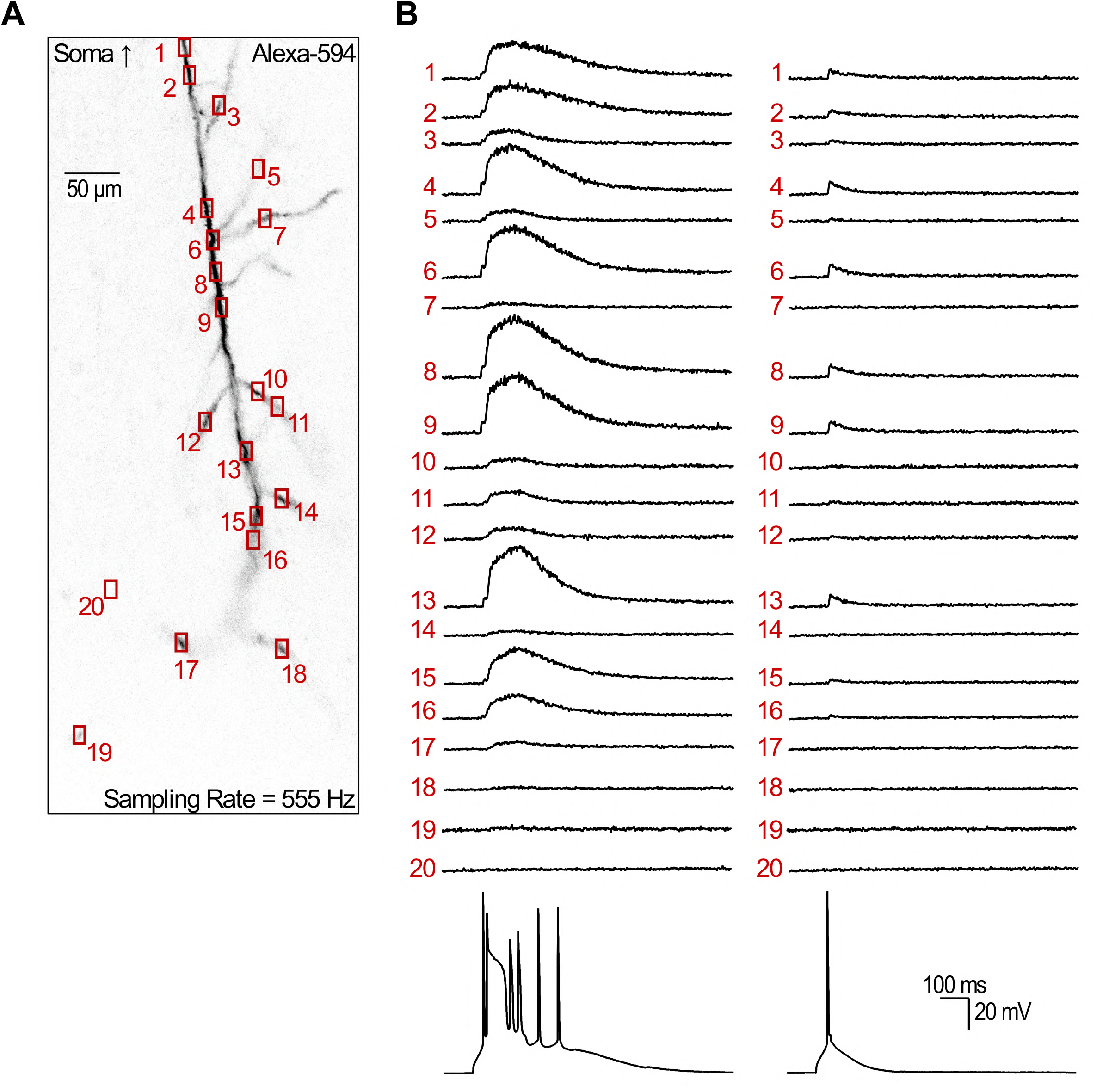
Random-access two-photon reveals the complex spatial profile of dendritic Ca^2+^ transients evoked by combined APs and plateaus or just APs. **A**, Single plane image showing the dendritic recording sites for that neuron. Random-access two-photon imaging allows for high-speed interrogation of the 20 recording sites at 555 Hz. ROI 20 is intentionally positioned away from the neurites. **B**, All traces associated with that neuron. Traces show the average of 3 trials.

